# Accurate and sensitive detection of microbial eukaryotes from whole metagenome shotgun sequencing

**DOI:** 10.1101/2020.07.22.216580

**Authors:** Abigail L. Lind, Katherine S. Pollard

## Abstract

**Background:** Microbial eukaryotes are found alongside bacteria and archaea in natural microbial systems, including host-associated microbiomes. While microbial eukaryotes are critical to these communities, they are challenging to study with shotgun sequencing techniques and are therefore often excluded.

**Results:** Here we present EukDetect, a bioinformatics method to identify eukaryotes in shotgun metagenomic sequencing data. Our approach uses a database of 521,824 universal marker genes from 241 conserved gene families, which we curated from 3,713 fungal, protist, non-vertebrate metazoan, and non-streptophyte archaeplastid genomes and transcriptomes. EukDetect has a broad taxonomic coverage of microbial eukaryotes, performs well on low-abundance and closely related species, and is resilient against bacterial contamination in eukaryotic genomes. Using EukDetect, we describe the spatial distribution of eukaryotes along the human gastrointestinal tract, showing that fungi and protists are present in the lumen and mucosa throughout the large intestine. We discover that there is a succession of eukaryotes that colonize the human gut during the first years of life, mirroring patterns of developmental succession observed in gut bacteria. By comparing DNA and RNA sequencing of paired samples from human stool, we find that many eukaryotes continue active transcription after passage through the gut, though some do not, suggesting they are dormant or nonviable. We analyze metagenomic data from the Baltic Sea and find that eukaryotes differ across locations and salinity gradients. Finally, we observe eukaryotes in *Arabidopsis* leaf samples, many of which are not identifiable from public protein databases.

**Conclusions:** EukDetect provides an automated and reliable way to characterize eukaryotes in shotgun sequencing datasets from diverse microbiomes. We demonstrate that it enables discoveries that would be missed or clouded by false positives with standard shotgun sequence analysis. EukDetect will greatly advance our understanding of how microbial eukaryotes contribute to microbiomes.

## Background

Eukaryotic microbes are ubiquitous in microbial systems, where they function as decomposers, predators, parasites, and producers [1]. Eukaryotic microbes inhabit virtually every environment on earth, from extremophiles found in geothermal vents, to endophytic fungi growing within the leaves of plants, to parasites inside the animal gut. Microbial eukaryotes have complex interactions with their hosts in both plant and animal associated microbiomes. For example, in plants, microbial eukaryotes assist with nutrient uptake and can protect against herbivory [2]. In animals, microbial eukaryotes in the gastrointestinal tract metabolize plant compounds [3]. Microbial eukaryotes can also cause disease in both plants and animals, as in the case of oomycete pathogens in plants or cryptosporidiosis in humans [4,5]. In humans, microbial eukaryotes interact in complex ways with the host immune system, and their depletion and low diversity in microbiomes from industrialized societies mirrors the industrialization-driven “extinction” seen for bacterial residents in the microbiome [6–8]. Outside of host associated environments, microbial eukaryotes are integral to the ecology of aquatic and soil ecosystems, where they are primary producers, partners in symbioses, decomposers, and predators [9,10].

Despite their importance, eukaryotic species are often not captured in studies of microbiomes [11]. By far the most frequently used metabarcoding strategy for studying host-associated and environmental microbiomes targets the 16S ribosomal RNA gene, which is found exclusively in bacteria and archaea, and therefore cannot be used to detect eukaryotes. Alternatively, amplicon sequencing of eukaryotic-specific (18S) or fungal-specific (ITS) ribosomal genes are effectively used in studies specifically targeting eukaryotes. Our understanding of the eukaryotic diversity in host-associated and environmental microbiomes is largely due to advances in eukaryotic-specific amplicon sequencing [12–15]. However, like all amplicon strategies they can suffer from limited taxonomic resolution and difficulty in distinguishing between closely related species [16,17].

Unlike amplicon strategies, whole metagenome sequencing captures fragments of DNA from the entire pool of species present in a microbiome, including eukaryotes. Whole metagenome sequencing data is increasingly common in microbiome studies, as these data can be used to assemble new genomes, disambiguate strains, and reveal presence and absence of genes and pathways [18]. As well as these methods work for identifying bacteria and archaea, multiple challenges remain to robustly and routinely identify eukaryotes in whole metagenome sequencing datasets. First, eukaryotic species are estimated to be present in many environments at lower abundances than bacteria, and comprise a much smaller fraction of the reads of a shotgun sequencing library than do bacteria [1,19]. Despite this limitation, interest in detecting rare taxa and rare variants in microbiomes has increased alongside falling sequencing costs, resulting in more and more microbiome deep sequencing datasets where eukaryotes could be detected and analyzed in a meaningful way.

Apart from sequencing depth, a second major barrier to detecting eukaryotes from whole metagenome sequencing is the availability of methodological tools. Research questions about eukaryotes in microbiomes using whole metagenome sequencing have primarily been addressed by using eukaryotic reference genomes, genes, or proteins for either *k*-mer matching or read mapping approaches [20,21]. However, databases of eukaryotic genomes and proteins have widespread contamination from bacterial sequences, and these methods therefore frequently misattribute bacterial reads to eukaryotic species [22,23]. Gene-based taxonomic profilers, such as Metaphlan3, have been developed to detect eukaryotic species, but these target a small subset of microbial eukaryotes (122 eukaryotic species as of the mpa_v30 release) [24,25].

To enable routine detection of eukaryotic species from any environment sequenced with whole metagenome sequencing, we have developed EukDetect, a bioinformatics approach that aligns metagenomic sequencing reads to a database of universally conserved eukaryotic marker genes. As microbial eukaryotes are not a monophyletic group, we have taken an inclusive approach and incorporated marker genes from 3,713 eukaryotes, including 596 protists, 2010 fungi, 146 non-streptophyte archaeplastids, and 961 non-vertebrate metazoans (many of which are microscopic or near-microscopic). This gene-based strategy avoids the pitfall of spurious bacterial contamination in eukaryotic genomes that has confounded other approaches. The EukDetect pipeline will enable different fields using whole metagenome sequencing to address emerging questions about microbial eukaryotes and their roles in human, other host-associated, and environmental microbiomes.

We apply the EukDetect pipeline to public human-associated, plant-associated, and aquatic microbiome datasets to make inferences about the roles of eukaryotes in these environments. We show that EukDetect’s marker gene approach greatly expands the number of detectable disease-relevant eukaryotic species in host-associated and environmental microbiomes.

## Results

### Bacterial sequences are ubiquitous in eukaryotic genomes

Eukaryotic reference genomes and proteomes have many sequences that are derived from bacteria, which have entered these genomes either spuriously through contamination during sequencing and assembly, or represent true biology of horizontal transfer from bacteria to eukaryotes [22,23]. In either case, these bacterial-derived sequences overwhelm the ability of either *k*-mer matching or read mapping-based approaches to detect eukaryotes from microbiome sequencing, as bacteria represent the majority of the sequencing library from many microbiomes. We demonstrate this problem by simulating paired-end sequence reads from the genomes of 971 common human gut microbiome bacteria, representing all major bacterial phyla in the human gut - including Bacteroidetes, Actinobacteria, Firmicutes, Proteobacteria, and Fusobacteria [26]. We aligned these reads against a database of 2,449 genomes of fungi, protists, and metazoans taken from NCBI GenBank. We simulated 2 million 126-bp paired-end reads per bacterial genome. Even with stringent read filtering (mapping quality > 30, aligned portion > 100 base pairs), a large number of simulated reads aligned to the eukaryotic database, from almost every individual bacterial genome (Figure 1A). In total, 112 bacteria had more than 1% of simulated reads aligning to the eukaryotic genome database, indicating that the potential for spurious alignment to eukaryotes from human microbiome bacteria is not limited to a few bacterial taxa (Figure S1). The majority (1,367 / 2,449) of the eukaryotic genomes contained at least 100 bp of genome sequence where bacterial reads spuriously aligned; 318 genomes had more than 1 Kb of bacterial sequence. These genomes came from all taxonomic groups tested (Figure 1C), indicating that bacterial contamination of eukaryotic genomes is widespread, and that simply removing a small number of contaminated genomes from an analysis will not be sufficient. As the majority of the sequencing library from many human microbiome samples is expected to be mostly comprised of bacteria [19], this spurious signal drowns out any true eukaryotic signal.

**Figure 1.**
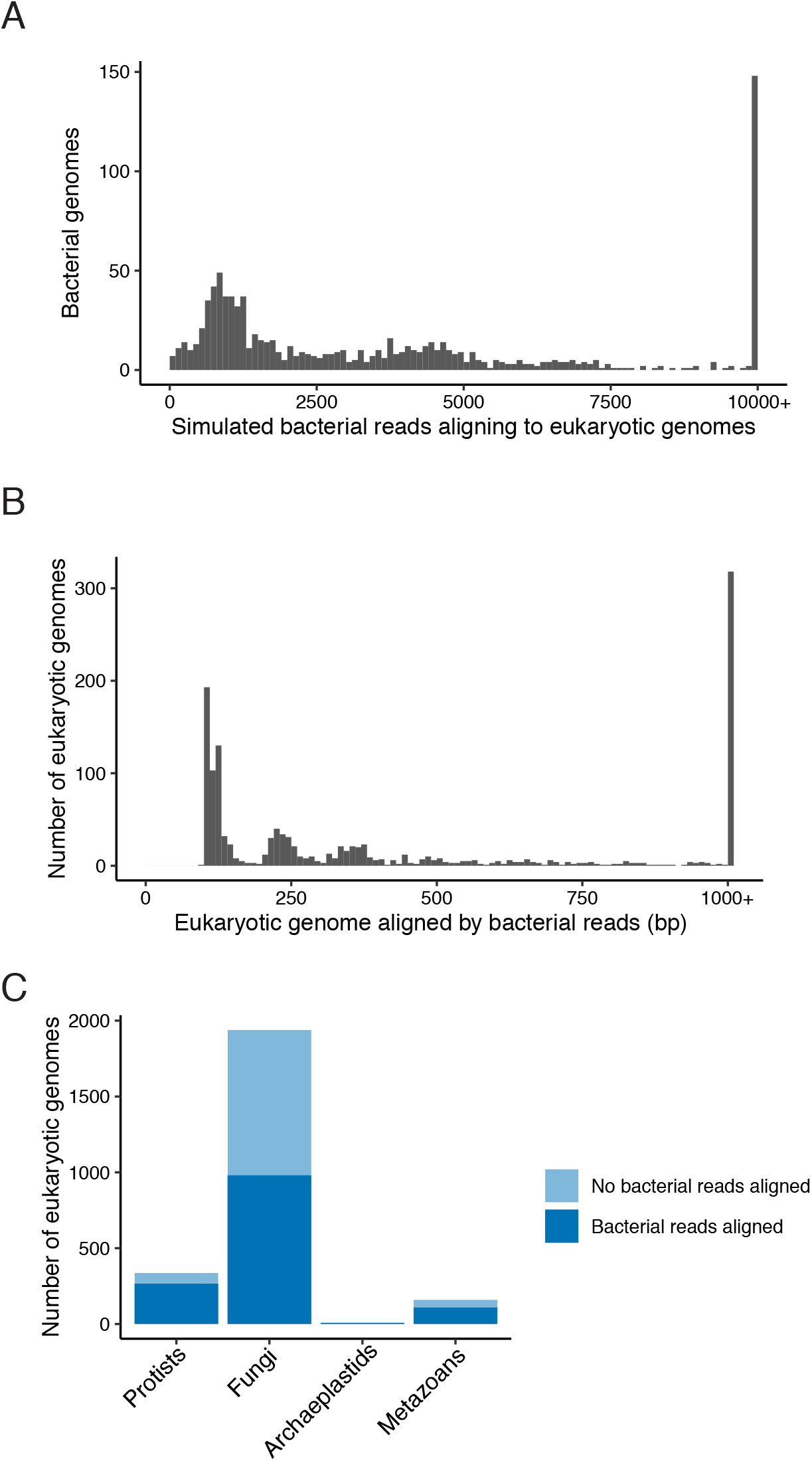
Human gut microbiome bacterial sequence reads are misattributed to eukaryotes. (A) Metagenomic sequencing reads were simulated from 971 species total from all major phyla in human stool (2 million reads per species) and aligned to all microbial eukaryotic genomes used to develop EukDetect. Even after stringent filtering (see Methods), many species have thousands of reads aligning to eukaryotic genomes, which would lead to false detection of eukaryotes in samples with only bacteria. (B) Amount of eukaryotic genome sequence aligned by simulated bacterial reads in 1,367 eukaryotic genomes. (C) Taxonomic distribution of eukaryotes in whole-genome database. Dark blue indicates eukaryotic genomes where bacterial reads aligned.

In practice, progress has been made in identifying eukaryotes from whole metagenome sequencing data by using alignment based approaches coupled with extensive masking, filtering, and manual curation [20,21,27]. However, these approaches still typically require extensive manual examination because of the incomplete nature of the databases used to mask. These manual processes are not scalable for analyzing the large amounts of sequencing data that are generated by microbiome sequencing studies and the rapidly growing number of eukaryotic genomes.

### Using universal marker genes to identify eukaryotes

To address the problem of identifying eukaryotic species in microbiomes from whole metagenome sequencing data, we created a database of marker genes that are uniquely eukaryotic in origin and uncontaminated with bacterial sequences. We chose genes that are ostensibly universally conserved to achieve the greatest identification power. We focused on universal eukaryotic genes rather than clade-, species-, or strain-specific genes because many microbial eukaryotes have variable size pangenomes and universal genes are less likely to be lost in a given lineage [28,29]. The chosen genes contain sufficient sequence variation to be informative about which eukaryotic species are present in sequencing data.

More than half of eukaryotic microbial species with sequenced genomes in NCBI Genbank do not have annotated genes or proteins. Therefore, we used the Benchmarking Universal Single Copy Orthologs (BUSCO) pipeline, which integrates sequence similarity searches and *de novo* gene prediction to determine the location of putative conserved eukaryotic genes in each genome [30]. Other databases of conserved eukaryotic genes exist, including PhyEco and the recently available EukCC and EukProt [31–33]. We found the BUSCO pipeline was advantageous, as BUSCO and the OrthoDB database have been benchmarked in multiple applications including gene predictor training, metagenomics, and phylogenetics [34,35].

We ran the BUSCO pipeline using the OrthoDB Eukaryote lineage [35] on 3,713 eukaryotic genomes and transcriptomes (2,010 fungi, 596 protists, 961 non-vertebrate metazoans, and 146 non-streptophyte archaeplastids) downloaded from NCBI GenBank or curated from other sources into the EukProt database, most of which do not have gene or protein predictions deposited in public databases [31] (Figure 2A). Marker gene sequences were then extensively quality filtered (see Methods). As part of the filtering process, we removed ~71,000 marker genes that were potentially bacterial in origin or contained regions similar to bacterial genomes, which comprised ~16% of the unfiltered database. Genes with greater than 99% sequence identity were combined and represent the most recent common ancestor of the species with collapsed genes. A final set of 521,824 marker genes from 214 BUSCO orthologous gene sets was selected, of which 500,508 genes correspond to individual species and the remainder to internal nodes (i.e., genera, families) in the taxonomy tree (Figure 2B, Table S3).

**Figure 2.**
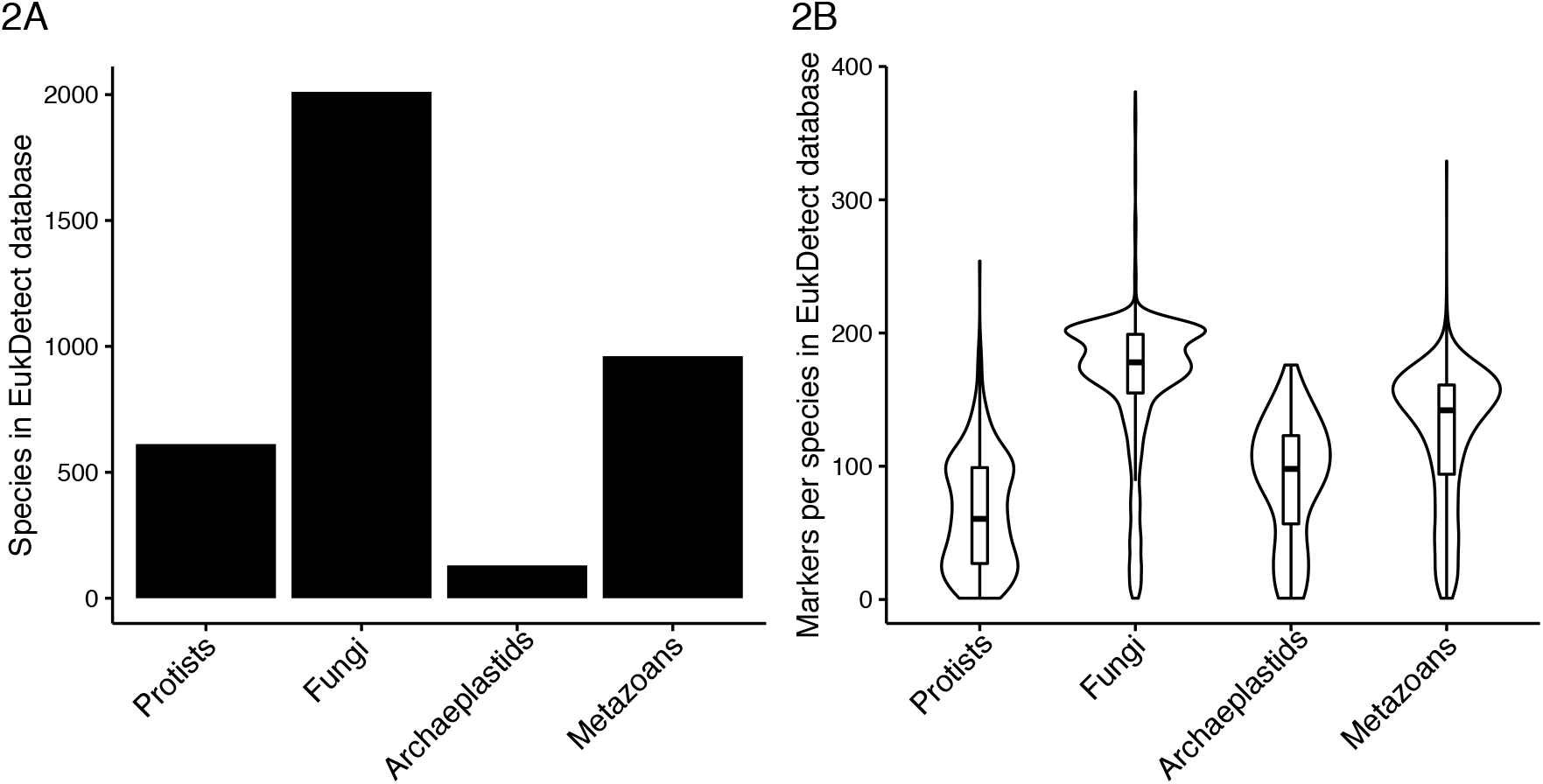
The EukDetect database comprises marker genes from protists, fungi, archaeplastids, and metazoans. (A) Taxonomic distribution of the species with transcriptomes and genomes included in the EukDetect database. (B) Total number of marker genes identified per species by taxonomic group.

Confirming that reads from bacteria would not spuriously align to this eukaryotic marker database, zero reads from our simulated bacterial sequencing data (Figure 1), or from reads simulated from ~55,000 human-associated and environmental metagenome assembled genomes (see Methods) align to this database. To further investigate the possibility that bacterial sequences may be present in marker genes, we analyzed 6,661 whole metagenome samples from 7 studies of human-associated, plant-associated, and aquatic microbiomes (see Methods), and compared the number of reads aligning to a given species to the median coverage of observed marker genes (Figure S2). We did not observe any cases where large numbers of reads align to small portions of marker genes, demonstrating that in these real datasets, bacterial contamination does not contribute to detected eukaryotic species.

Despite the BUSCO universal single-copy nomenclature, the EukDetect database genes are present at variable levels in different species (Figure 2B). We uncovered a number of patterns that explain why certain taxonomic groups have differing numbers of marker genes. First, BUSCOs may be absent because the genomes for a species are incomplete. Second, many of the species present in our dataset, including microsporidia and many of the pathogenic protists, function as parasites. As parasites rely on their host for many essential functions, they frequently experience gene loss and genome reduction [36], decreasing the representation of BUSCO genes. It is also possible that the BUSCO pipeline does not identify all marker genes in some clades with few representative genomes. Conversely, some clades have high rates of duplication for these typically single-copy genes. These primarily occur in fungi; studies of fungal genomes have demonstrated that hybridization and whole genome duplication occurs frequently across the fungal tree of life [37,38].

As the representation of each marker gene differs across taxonomic groups, we identified a subset of 50 marker genes that are most frequently found across all eukaryotic taxa (Table S3). This core set of genes has the potential to be used for strain identification, abundance estimation, and phylogenetics, as has been demonstrated for universal genes in bacteria [33].

### EukDetect: a pipeline to accurately identify eukaryotes using universal marker genes

We incorporated the complete database of 214 marker genes into EukDetect, a bioinformatics pipeline (https://github.com/allind/EukDetect) that first aligns metagenomic sequencing reads to the marker gene database, and filters each read based on alignment quality, sequence complexity, and alignment length. Sequencing reads can be derived from DNA or from RNA sequencing. EukDetect then removes duplicate sequences, calculates the coverage and percent identity of detected marker genes, and reports these results in a taxonomy-informed way. When more than one species per genus is detected, EukDetect determines whether any of these species are likely to be the result of off-target alignment, and reports these hits separately (see Methods for more detail). Optionally, results can be obtained for only the 50 most universal marker genes.

### EukDetect is sensitive and accurate even at low sequence coverage

To determine the performance of EukDetect, we simulated reads from representative species found in host associated and environmental microbiomes, including one human-associated fungus (*Candida albicans*) and one soil fungus (*Trichoderma harzianum*), one human-associated protist (*Entameoba dispar*) and one environmental protist (the widely distributed ocean haptophyte *Emiliania huxleyi*), and one human-associated helminth (*Schistosoma mansoni*) and one plant pathogenic nematode (*Globodera rostochiensis*). These species represent different groups of the eukaryotic tree of life, and are variable in their genome size and representation in the EukDetect marker gene database. The number of reads aligning to the database and the number of observed marker genes vary based on the total number of marker genes per organism and the overall size of its genome (Figure 3). We simulated metagenomes with sequencing coverage across a range of values for each species, and performed 10 simulations for each species. We found that EukDetect is highly sensitive and aligns at least one read to 80% or more of all marker genes per species at low simulated genome coverages between 0.25x or 0.5x (Figure 3A).

**Figure 3.**
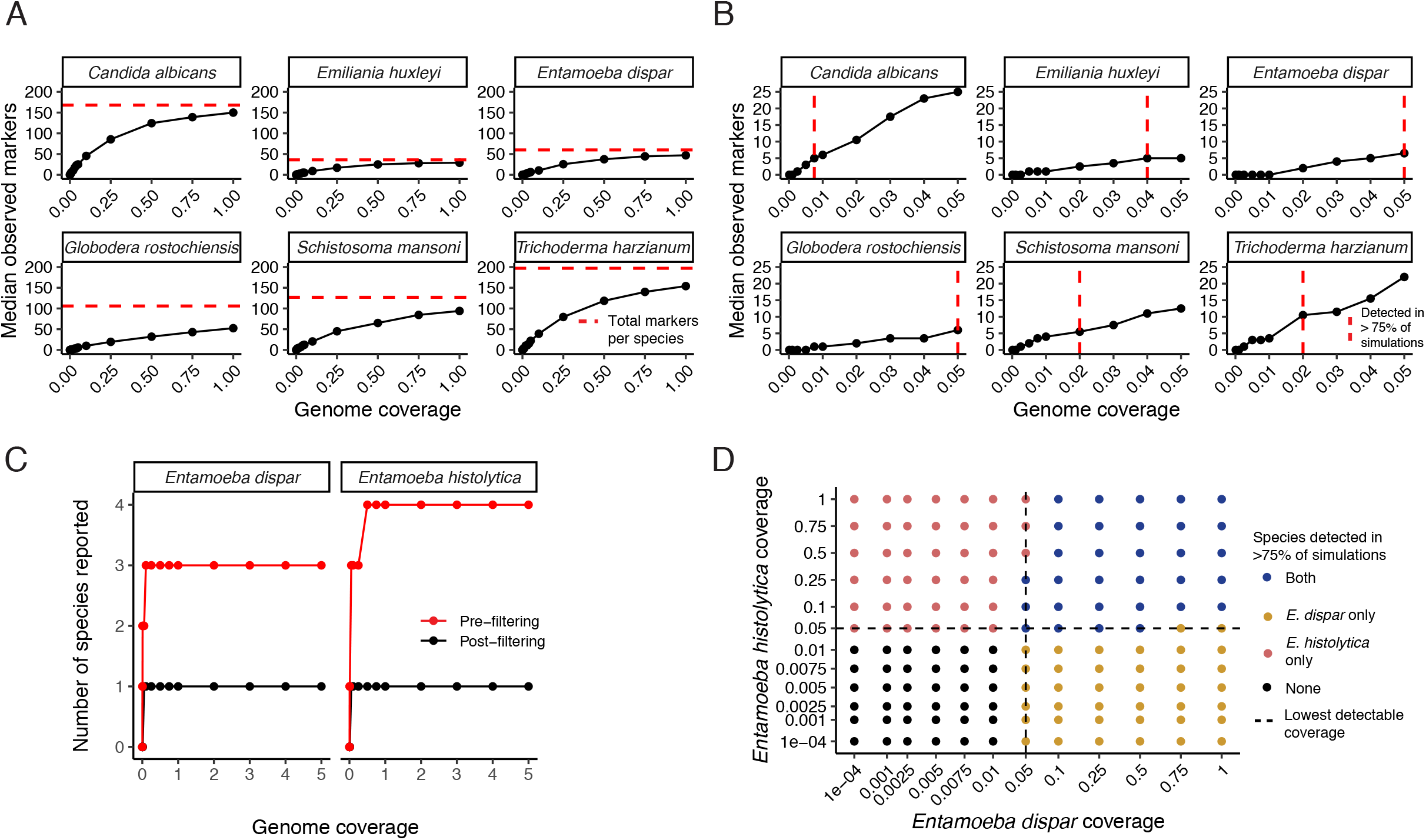
EukDetect is sensitive and accurate for yeasts, protists, and worms at low sequence coverage. (A) Number of marker genes with at least one aligned read per species up to 1x genome coverage. Horizontal red line indicates the total number of marker genes per species (i.e. the best possible performance). (B) Number of marker genes with at least one aligned read per species up to 0.05x genome coverage. Vertical red line indicates a detection cutoff where 4 or more reads align to 2 or more markers in 8 out of 10 simulations or more. (C) The number of species reported by EukDetect for two closely related Entamoeba species before and after minimum read count and off-target alignment filtering. One species is the correct result. Simulated genome coverages are the same as in panel D. (D) EukDetect performance on simulated sequencing data from mixtures of two closely related Entamoeba species at different genome coverages. Dot colors indicate which species were detected in 8 out of 10 simulations or more. Dashed line indicates the lowest detectable coverage possible by EukDetect when run on each species individually. Axes not to scale.

Using a detection limit cutoff requiring at least 4 unique reads aligning to 2 or more marker genes, *Candida albicans* is detectable in 8 or more out of 10 simulations at 0.005x coverage (Figure 3B). *Schistosoma mansoni* and *Trichoderma harzianum* are both detected in 8 or more out of 10 simulations at 0.01x coverage. *Emiliania huxleyi* and *Globodera rostochiensis* are both detected in 8 or more out of 10 simulations at 0.05x coverage, and *Acanthamoeba castellanii* is detected in 8 or more out of 10 simulations at 0.1x coverage.

Off-target alignment happens when reads from one species erroneously align to another, typically closely related, species. This can occur due to errors in sequencing or due to high local similarities between gene sequences. To account for this error, when more than one species from a genus is detected in a sample, EukDetect examines the reads aligning to each species and determines whether any detected species are likely to have originated from off-target alignment. To test this approach, we simulated reads from the genomes of two closely related species found in human microbiomes, the amoebozoan parasite *Entamoeba histolytica* and the amoebozoan commensal *Entamoeba dispar*. These species are morphologically identical, and their genomes have an average nucleotide identity of 90% (calculated with fastANI [39]). While simulated reads from both genomes do align to other closely related *Entamoeba* species, EukDetect accurately marks these as off-target hits and does not report them in the primary results files (Figure 3C).

To determine whether EukDetect can accurately distinguish true mixtures of very closely related species, we simulated combinations of both *Entamoeba* species with genome coverage from 0.0001x coverage up to 1x coverage and determined when EukDetect results report both, either one, or neither species (Figure 3D). In general, EukDetect accurately detected each of the species when present at above the detection limit we determined from simulations of the single species alone (0.01x coverage for *Entamoeba histolytica* and 0.0075x coverage for *Entamoeba dispar*) in 8 or more out of 10 simulations. However, in several edge cases where one species was present at or near its detection limit and the other species was much more abundant, EukDetect was not able to differentiate the lower abundance species from possible off-target alignments and did not report them in the final output. As pre-filtered results are reported as a secondary output by EukDetect, users can examine these files for evidence of a second species in cases where two closely related eukaryotic species are expected in a sample.

One additional scenario that may produce reads aligning to multiple species within a genus is if a sample contains a species that is not in the EukDetect marker gene database but is within a genus that has representatives in the database. The most conservative option is to consider the most recent common ancestor of the closely related species, which usually resolves at the genus level. This information is provided by EukDetect (Figure S3).

### Vignettes of microbial eukaryotes in microbiomes

To demonstrate how EukDetect can be used to understand microbiome eukaryotes, we investigated several biological questions about eukaryotes in microbiomes using publicly available datasets from human- and plant-associated microbiomes.

#### The spatial distribution of eukaryotic species in the gut

While microbes are found throughout the human digestive tract, studies of the gut microbiome often examine microbes in stool samples, which cannot provide information about their spatial distribution [40]. Analyses of human and mouse gut microbiota have shown differences in the bacteria that colonize the lumen and the mucosa of the GI tract, as well as major differences between the upper GI tract, the small intestine, and the large intestine, related to the ecology of each of these environments. Understanding the spatial organization of microbes in the gut is critical to dissecting how these microbes interact with the host and contribute to host phenotypes.

To determine the spatial distribution of eukaryotic species in the human gut, we analyzed data from two studies that examined probiotic strain engraftment in the human gut. These studies used healthy adult humans recruited in Israel, and collected stool samples along with biopsies performed by endoscopy from 18 different body sites along the upper GI tract, small intestine, and large intestine [41,42]. A total of 1,613 samples from 88 individuals were sequenced with whole metagenome sequencing.

In total, eukaryotes were detected in 325 of 1,613 samples with the EukDetect pipeline (Table S4). The most commonly observed eukaryotic species were subtypes of the protist *Blastocystis* (236 samples), the yeast *Saccharomyces cerevisiae* (66 samples), the protist *Entamoeba dispar* (17 samples), the yeast *Candida albicans* (9 samples), and the yeast *Cyberlindnera jadinii* (6 samples). Additional fungal species were observed in fewer samples, and included the yeast *Malassezia restricta,* a number of yeasts in the *Saccharomycete* class including *Candida tropicalis* and *Debaromyces hansenii*, and the saprophytic fungus *Penicillium roqueforti*.

The prevalence and types of eukaryotes detected varied along the GI tract. In the large intestine and the terminal ileum, microbial eukaryotes were detected at all sites, both mucosal and lumen-derived. However, they were mostly not detected in the small intestine or upper GI tract (Figure 4). One exception to this is a sample from the gastric antrum mucosa which contained sequences assigned to the fungus *Malassezia restricta. Blastocystis* species were present in all large intestine and terminal ileum samples, while fungi were present in all large intestine samples and the lumen of the terminal ileum. The protist *Entamoeba dispar* was detected almost exclusively in stool samples; only one biopsy-derived sample from the descending colon lumen contained sequences assigned to an *Entamoeba* species. The saprophytic fungus *Penicillium roqueforti*, which is used in food production and is likely to be allochthonous in the gut [43], was only detected in stool samples.

**Figure 4.**
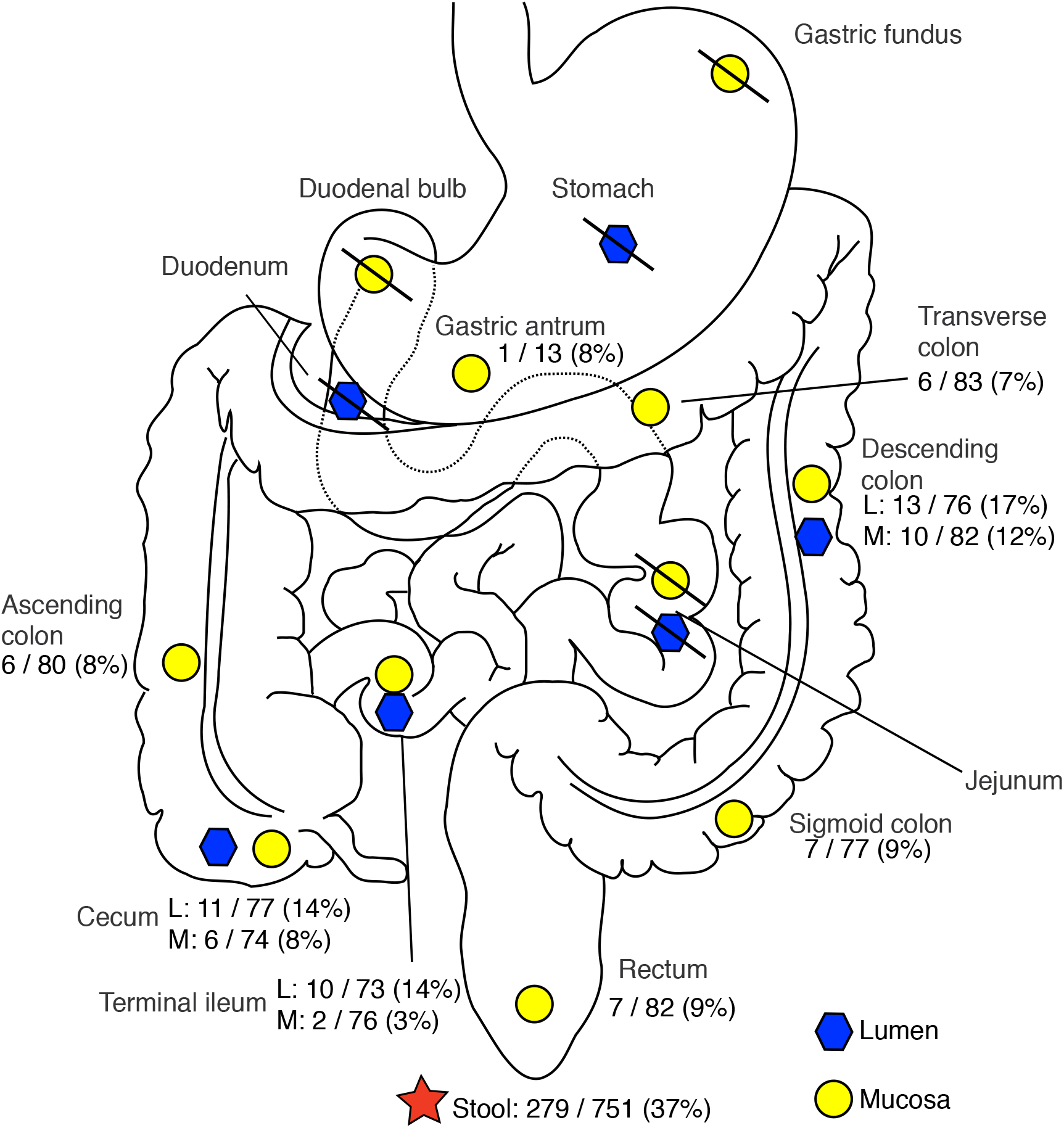
Distribution of eukaryotic species in the gastrointestinal tract taken from biopsies. Eukaryotes were detected at all sites in the large intestine and in the terminal ileum, in both lumen and mucosal samples. One biopsy of gastric antrum mucosa in the stomach contained a *Malassezia* yeast. Slashes indicate no eukaryotes detected in any samples from that site. See Figures S3 and S4 for locations of *Blastocystis* subtypes and locations of fungi.

Our detection of microbial eukaryotes in mucosal sites suggests that they may be closely associated with host cells and not just transiently passing through the GI tract. We detected both protists and fungi (*Blastocystis, Malassezia*, and *Saccharomyces cerevisiae*) in mucosal as well as lumen samples. These findings are consistent with previous studies of *Blastocystis-infected* mice that found *Blastocystis* in both the lumen and the mucosa of the large intestine [44]. Taken together, our findings indicate that certain eukaryotic species can directly interact with the host, similar to mucosal bacteria [40].

#### A succession of eukaryotic microbes in the infant gut

The gut bacterial microbiome undergoes dramatic changes during the first years of life [45]. We sought to determine whether eukaryotic members of the gut microbiome also change over the first years of life. To do so, we examined longitudinal whole metagenome sequencing data from the three-country DIABIMMUNE cohort, where stool samples were taken from infants in Russia, Estonia, and Latvia during the first 1200 days of life [46]. In total, we analyzed 791 samples from 213 individuals.

Microbial eukaryotes are fairly commonly found in stool in the first few years of life. We detected reads assigned to a eukaryotic species from 108 samples taken from 68 individuals (Table S5). The most frequently observed species were *Candida parapsilosis* (31 samples), *Saccharomyces cerevisiae* (29 samples), various subtypes of the protist *Blastocystis* (18 samples), *Malassezia* species (14 samples), *Candida albicans* (8 samples), and *Candida lusitaniae* (8 samples). These results are consistent with previous reports that *Candida* species are prevalent in the neonatal gut [47,48]. Less frequently observed species were primarily yeasts in the *Saccharomycete* class; the coccidian parasite *Cryptosporidium meleagridis* was detected in one sample.

These species do not occur uniformly across time. When we examined covariates associated with different eukaryotic species, we found a strong association with age. The median age at collection of the samples analyzed with no eukaryotic species was 450 days, but *Blastocystis* protists and *Saccharomycetaceae* fungi were primarily observed among older infants (median 690 days and 565 days, respectively) (Figure 5A). Fungi in the *Debaromycetaceae* family were observed among younger infants (median observation 312 days). Samples containing fungi in the *Malasseziaceae* family were older than *Debaromycetaceae* samples (median observation 368 days), though younger than the non-eukaryotic samples. To determine whether these trends were statistically significant, we compared the mean ages of two filtered groups: individuals with no observed eukaryotes and individuals where only one eukaryotic family was observed (Figure 5B). We found that *Saccharomycetaceae* fungi and *Blastocystis* protists were detected in significantly older children (Wilcoxon rank-sum p=0.0012 and p=2.6e-06, respectively). In contrast, *Debaromycetaceae* fungi were found in significantly younger infants (p=0.035). As only three samples containing *Malasseziaceae* came from individuals where no other eukaryotic families were detected, we did not analyze this family.

**Figure 5.**
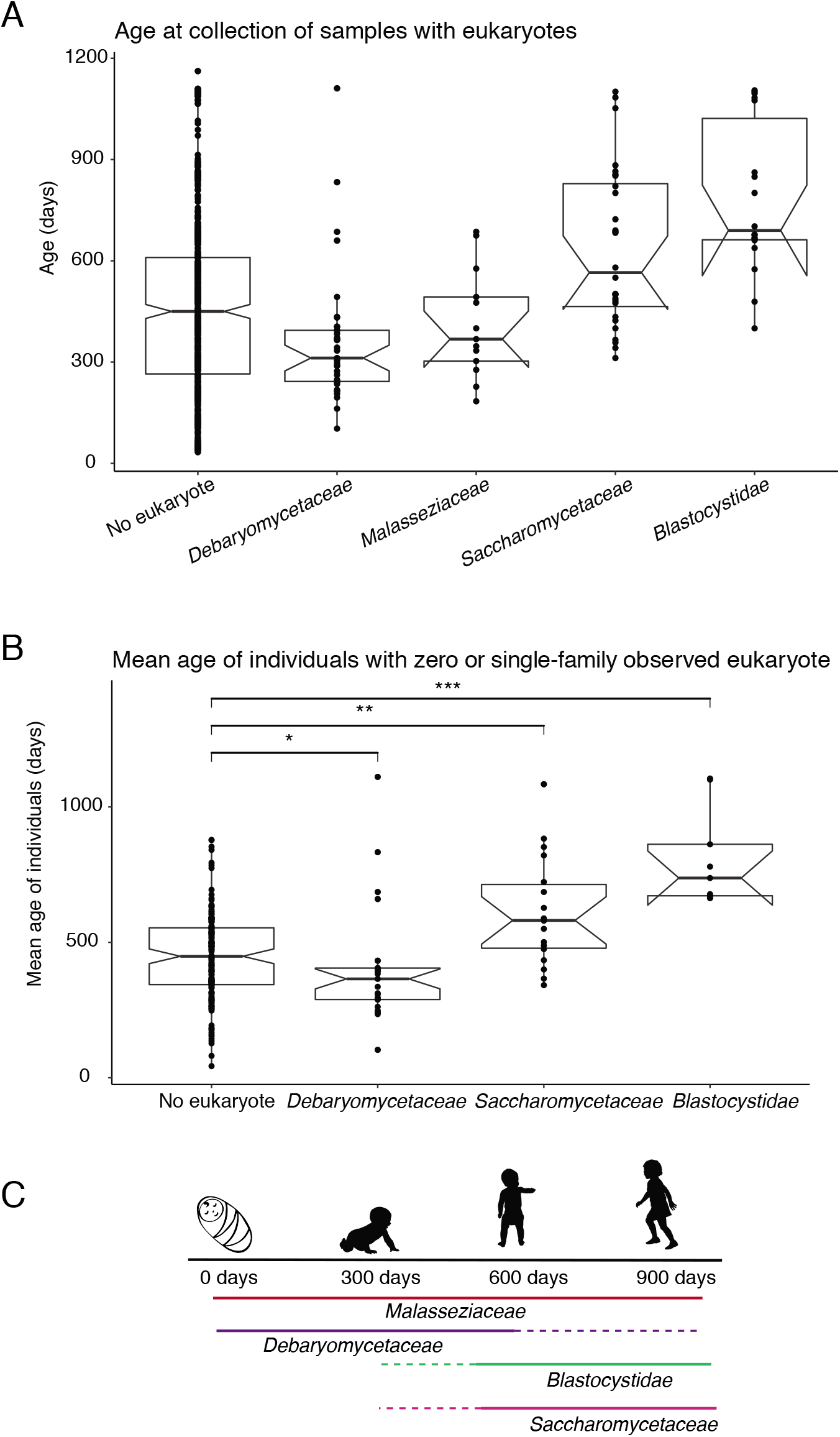
Changes in eukaryotic gut microbes during the first years of life. (A) Age at collection in the DIABIMMUNE three-country cohort for samples with no eukaryote or with any of the four most frequently observed eukaryotic families. (B) The mean age at collection of samples from individuals with no observed eukaryotes compared to the mean age at collection of individuals where one of three eukaryotic families were detected. Individuals where more than one eukaryotic family was detected are excluded. *Malasseziaceae* is excluded due to low sample size. Group comparisons were performed with an unpaired Wilcoxon rank-sum test. *, p < 0.05; **, p < 0.01; ***, p < 0.001. (C) Model of eukaryotic succession in the first years of life. *Debaryomycetaceae* species predominate during the first two years of life, but are also detected later. *Blastocystidae* species and *Saccharomycetaceae* species predominate after the first two years of life, though they are detected as early as the second year of life. *Malasseziaceae* species do not change over time.

These findings suggest that, as observed with bacteria, the eukaryotic species that colonize the gastrointestinal tract of children change during early life [45]. Our results support a model of eukaryotic succession in the infant gut, where *Debaromycetaceae* fungi, notably *Candida parapsilosis* in these data, dominate the eukaryotic fraction of the infant gut during the first year of life, and *Blastocystis* and *Saccharomyces* fungi, which are commonly observed in the gut microbiomes of adults, rise to higher prevalence in the gut in the second year of life and later (Figure 5C). Altogether these results expand the picture of eukaryotic diversity in the human early life GI tract.

#### Differences in RNA and DNA detection of eukaryotic species suggests differential transcriptional activity in the gut

While most sequencing-based analyses of microbiomes use DNA, microbiome transcriptomics can reveal the gene expression of microbes in a microbiome, which has the potential to shed light on function. RNA sequencing of microbiomes can also be used to distinguish dormant or dead cells from active and growing populations. We sought to determine whether eukaryotic species are detectable from microbiome transcriptomics, and how these results compare to whole metagenome DNA sequencing.

We leveraged data from the Inflammatory Bowel Disease (IBD) Multi’omics Data generated by the Integrative Human Microbiome Project (IHMP), which generated RNA-Seq, whole-genome sequencing, viral metagenomic sequencing, and 16S amplicon sequencing from stool samples of individuals with Crohn’s disease, ulcerative colitis, and no disease (IBDMDB; http://ibdmdb.org) [49]. We analyzed samples with paired whole metagenome sequencing and RNA sequencing. In the IHMP-IBD dataset, there were 742 sets of paired DNA and RNA sequencing from a sample from 104 individuals, 50 of whom were diagnosed with Crohn’s disease, 28 of whom were diagnosed with ulcerative colitis, and 19 with no with IBD. Samples were collected longitudinally, and the median number of paired samples per individual was seven.

Microbial eukaryotes were prevalent in this dataset. EukDetect identified eukaryotic species in 409 / 742 RNA-sequenced samples and 398 / 742 DNA-sequenced samples. The most commonly detected eukaryotic species were *Saccharomyces cerevisiae, Malassezia* species, *Blastocystis* species, and the yeast *Cyberlindnera jadinii* (Table S6). Eukaryotic species that were detected more rarely were primarily yeasts from the *Saccharomycete* class. We examined whether eukaryotic species were present predominantly in DNA sequencing, RNA sequencing, or both, and found different patterns across different families of species (Figure 6). Of the 585 samples where we detected fungi in the *Saccharomycetaceae* family, 314 of those samples were detected from the RNA-sequencing alone and not from the DNA. A further 178 samples had detectable *Saccharomycetaceae* fungi in both the DNA and the RNA sequencing, while 17 samples only had detectable *Saccharomycetaceae* fungi in the DNA sequencing. Fungi in the *Malasseziaceae* family were only detected in DNA sequencing (115 samples total), and the bulk of *Cyberlindnera jadinii* and *Debaryomycetaceae* fungi (*Candida albicans* and *Candida tropicalis*) were detected in DNA sequencing alone. In contrast, *Blastocystis* protists were mostly detected both from RNA and from DNA, and *Pichiaceae* fungi (*Pichia* and *Brettanomyces* yeasts) were detected in both RNA and DNA sequencing, DNA sequencing alone, and in a small number of samples in RNA-seq alone.

**Figure 6.**
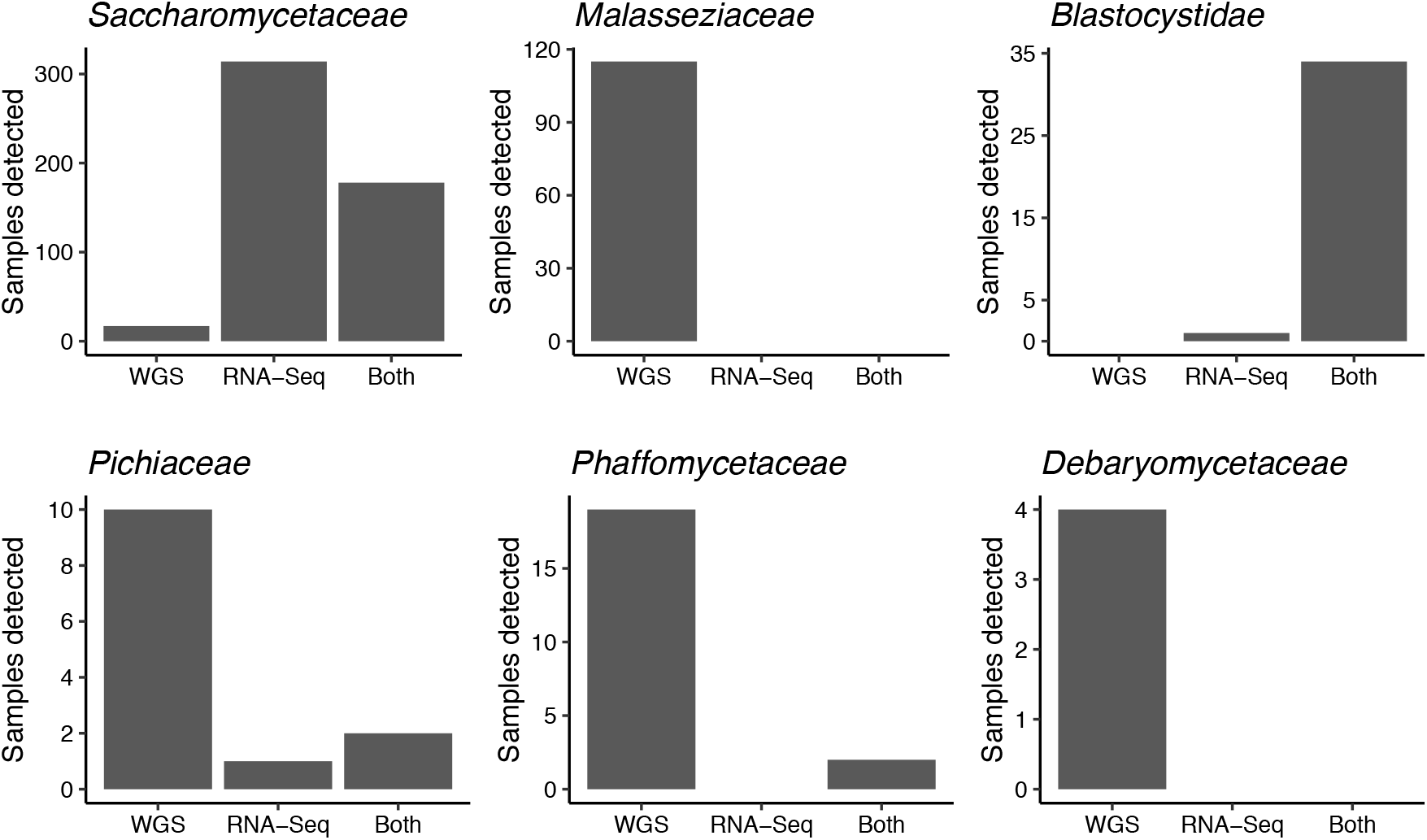
Detection of eukaryotes from paired DNA and RNA sequenced samples from the IHMP IBD cohort. Plots depict the most commonly detected eukaryotic families, and whether a given family was detected in the DNA sequencing alone, the RNA sequencing alone, or from both the RNA and the DNA sequencing from a sample. Some samples shown here come from the same individual sampled at different time points.

From these findings, we can infer information about the abundance and possible functions of these microbial eukaryotes. *Blastocystis* is the most frequently observed gut protist in the human GI tract in industrialized nations [50], and its high relative abundance is reflected in the fact that it is detected most frequently from both RNA and DNA sequencing. The much greater detection of *Saccharomycetaceae* fungi from RNA than from DNA suggests that these cells are transcriptionally active, and that while the absolute cell counts of these fungi may not be detectable from DNA sequencing, they are actively transcribing genes and therefore may impact the ecology of the gut microbiome. In contrast, fungi in the *Malasseziaceae* family are found at high relative abundances on human skin, and while they have been suggested to play functional roles in the gut [51–53], the data from this cohort suggest that the *Malasseziaceae* fungi are rarely transcriptionally active by the time they are passed in stool. Fungi in the *Phaffomycetaceae, Pichiaceae*, and *Debaromycetaeae* families likely represent a middle ground, where some cells are transcriptionally active and detected from RNA-sequencing, but others are not active in stool. These results suggest that yeasts in the *Saccharomycete* clade survive passage through the GI tract and may contribute functionally to the gut microbiome.

#### Eukaryotes in the Baltic Sea differ across environments and salinity gradients

As the EukDetect marker database contains eukaryotic species from all environments, it can be used to detect eukaryotes outside of the animal gut microbiome. As one example, we applied it to whole metagenome sequencing from the Baltic Sea. The Baltic Sea is one of the world’s largest brackish bodies of water, and it contains multiple geochemical gradients across its surface and depths including salinity and oxygen gradients [54]. Influx of fresh water into the Baltic Sea creates a halocline, which results in low mixing between upper oxygenated and lower anoxic waters. This stratification results in an intermediate layer with a strong vertical redox gradient, which is referred to as the redoxcline.

We examined 80 whole metagenome sequenced samples from the Baltic Sea sampled across geographic location, depth, and geochemical characteristics for eukaryotic species using EukDetect. Of these samples, 37 were obtained at the Linneaus Microbial Observatory near Öland, Sweden (referred to as LMO samples) [55]. An additional 30 samples were taken from 9 different geographic locations across the Baltic Sea at different depths (referred to as transect samples), and 14 samples collected across different zones of the redoxcline (referred to as redox samples).

EukDetect identified one or more eukaryotic species in 75 or these 80 samples (Table S7, Figure S6). All samples from the LMO and the Transect sets contained eukaryotes; 9 of the 14 Redox samples contained eukaryotes (Figure S6). The four most frequently observed species were chloroplastids (Figure 7A); in total, 70 samples contained at least one chloroplastid species, with species in the Mamiellophyceae class being most widespread (Figure S7). Protists were also frequently detected; stramenopiles, in particular diatoms (Figure S7), were detected in 42 samples, while haptophytes and alveolates were detected in 33 and 29 samples, respectively (Figure 7A).

**Figure 7.**
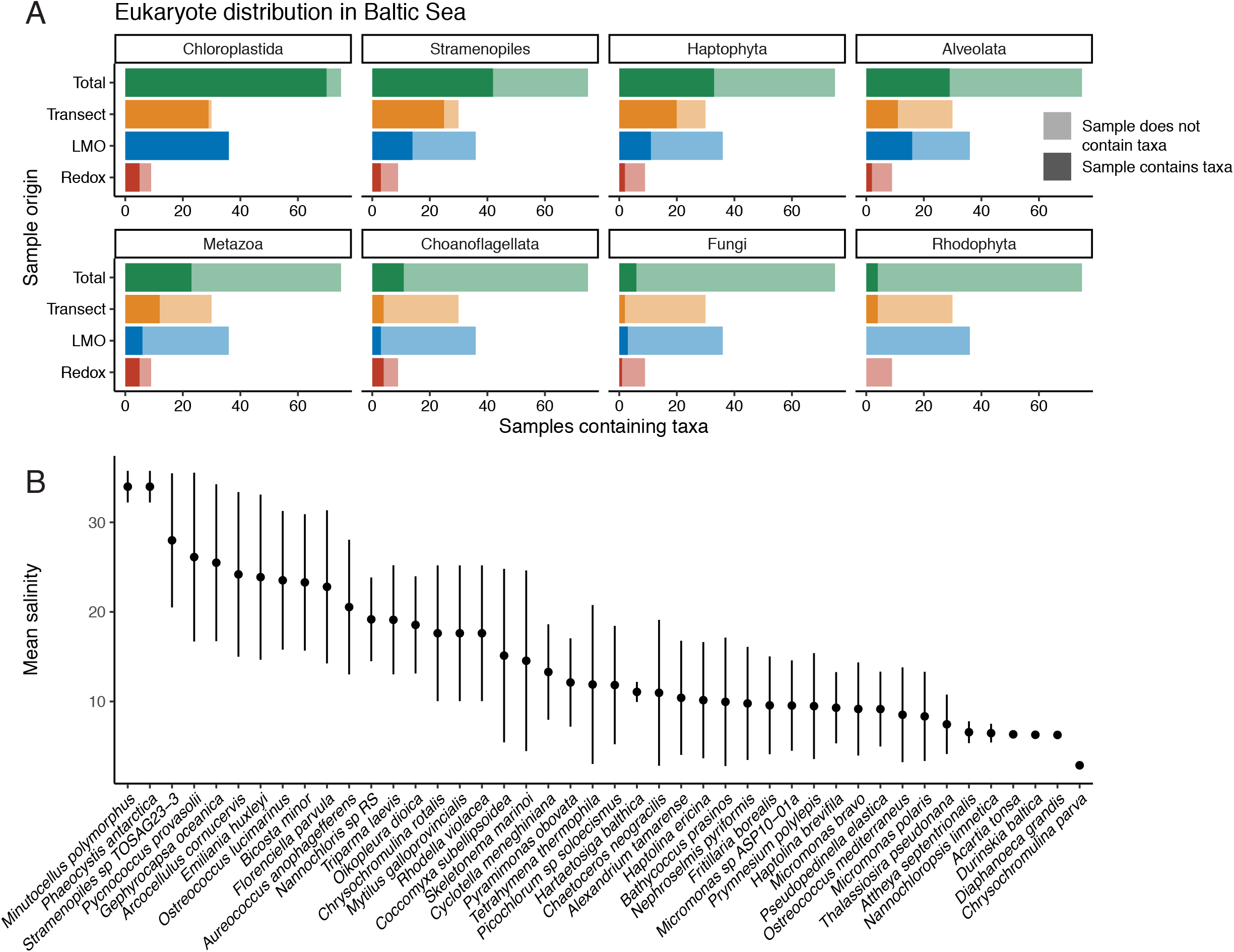
Eukaryotes in the Baltic Sea differ across environments and salinity gradients. (A) Counts of observed eukaryotic groups detected in the Baltic Sea across different environment. LMO samples were obtained at the Linneaus Microbial Observatory near Öland, Sweden. Transect samples were obtained from 9 different geographic locations across the Baltic Sea. Redox samples were obtained from the redoxcline. Dark colored bars indicate the number of samples from a given environment that contain at least one species belonging to the eukaryotic subgroup. Light colored bars indicate the number of samples that do not contain a given eukaryotic subgroup. (B) The mean salinity of samples where eukaryotic species were detected. Bars indicate 1 standard deviation. Only species detected in 3 or more samples were included.

Eukaryotes detected in the redoxcline differed from those seen in LMO and transect samples. The most frequently observed species in these samples was the choanoflagellate *Hartaetosiga balthica* (prev. *Codosiga balthica*), which was detected in four redox samples (Table S7). This choanoflagellate has been previously described to inhabit hypoxic waters of the Baltic Sea, and it possesses derived mitochondria believed to reflect an adaptation to hypoxia [56]. Notably, this choanoflagellate is the only observed eukaryotic species in samples with zero dissolved oxygen. Two additional species exclusively detected in the redoxcline are an ascomycete fungus *Lecanicillium* sp. LEC01, which was originally isolated from jet fuel [57], and the bicosoecid marine flagellate *Cafeteria roenbergensis* (Table S7).

Salinity is a major driver in the composition of bacterial communities in the Baltic Sea [58]. To determine how eukaryotes vary across the salinity gradients in the Baltic Sea, we examined the mean and variance of the salinity of eukaryotes detected in three or more samples (Figure 7B). Some eukaryotic species are found across broad ranges of salinities, while others appear to prefer narrower ranges. Eukaryotes found across a broad range of salinities may be euryhaline, or alternatively, different populations may undergo local adaptation to salinity differences. The eukaryote found across the broadest range of salinities, the diatom *Skeletonema marinoi*, was shown previously to undergo local adaptation to salinity in the Baltic Sea [59]. Further, one eukaryote found over a broad range of salinities, *Tetrahymena thermophila*, readily undergoes local adaptation in response to temperature gradients in experimental evolution experiments [60]. We hypothesize that similar processes are occurring in response to salinity gradients in the Baltic Sea.

#### Eukaryotes in the plant leaf microbiome

EukDetect can also be used to detect eukaryotes in non-animal host associated microbiomes. To demonstrate this, we analyzed 1,394 samples taken from 275 wild *Arabidopsis thaliana* leaves [61]. From these samples, we detect 37 different eukaryotic species and 25 different eukaryotic families (Table S8). We found pathogenic *Peronospora* oomycetes in 374 samples and gall forming *Protomyces* fungi in 311 samples. Other frequently observed eukaryotes in these samples are *Dothideomycete* fungi in 101 samples, *Sordariomycete* fungi in 29 samples, *Agaricomycete* fungi in 26 samples, and *Tremellomycete* fungi in 10 samples*. Malassezia* fungi are also detected in 14 samples, though we hypothesize that these are contaminants from human skin. Many of the most commonly detected microbial eukaryotic species in these samples, including the *Arabidopsis*-isolated yeast *Protomyces* sp. C29 in 311 samples and the epiphytic yeast *Dioszegia crocea* in 10 samples, do not have annotated genes associated with their genomes and have few to no gene or protein sequences in public databases. These taxa and others would not be identifiable by aligning reads to existing gene or protein databases. These results demonstrate that EukDetect can be used to detect microbial eukaryotes in non-human host-associated microbiomes, and reveal more information than aligning reads to gene or protein databases.

## Discussion

Here we have presented EukDetect, an analysis pipeline that leverages a database of conserved eukaryotic genes to identify eukaryotes present in metagenomic sequencing. By using conserved eukaryotic genes, this pipeline avoids inaccurately classifying sequences based on the widespread bacterial contamination of eukaryotic genomes (Figure 1) [22,23]. The EukDetect pipeline is sensitive and can detect fungi, protists, non-streptophyte archaeplasts, and non-vertebrate metazoans present at low sequencing coverage in metagenomic sequencing data.

We apply the EukDetect pipeline to public datasets from the human gut microbiome, the plant leaf microbiome, and an aquatic microbiome, detecting eukaryotes uniquely present in each environment. We find that fungi and protists are present at all sites within the lower GI tract of adults, and that the eukaryotic composition of the gut microbiome changes during the first years of life. Using paired DNA and RNA sequencing from the iHMP IBDMBD project, we demonstrate the utility of EukDetect on RNA data and show that some eukaryotes are differentially detectable in DNA and RNA sequencing, suggesting that some eukaryotic cells are dormant or dead in the GI tract, while others are actively transcribing genes. We find that salinity gradients impact eukaryote distribution in the Baltic Sea. Finally, we find oomycetes and fungi in the *Arabidopsis thaliana* leaf microbiome, many of which would not have been detectable using existing protein databases.

One important limitation of our approach is that only eukaryotic species with sequenced genomes or that have close relatives with a sequenced genome can be detected by the EukDetect pipeline. Due to taxonomic gaps in sequenced genomes, the EukDetect database does not cover the full diversity of the eukaryotic tree of life. We focused our applications on environments that have been studied relatively well. But nonetheless some eukaryotes that are known to live in human GI tracts, such as *Dientamoeba fragilis* [62], have not been sequenced and would have been missed if they were present in the samples we analyzed. Improvements in single-cell sequencing [63] and in metagenomic assembly of eukaryotes [64] will increase the representation of uncultured eukaryotic microbes in genome databases. Because EukDetect uses universal genes, it will be straightforward to expand its database as more genomes are sequenced.

EukDetect enables the use of whole metagenome sequencing data, which is frequently generated in microbiome studies for many different purposes, for routine detection of eukaryotes. Major limitations of this approach are that eukaryotic species are often present at lower abundances than bacterial species and thus sometimes excluded from sequencing libraries, and that EukDetect is limited to species with sequenced genomes or transcriptomes. Marker gene sequencing, such as 18S for all eukaryotes, or ITS for fungi, is agnostic to whether a species has a deposited genome or transcriptome, and can detect lower-abundance taxa than can whole metagenome sequencing. However, unlike whole metagenome sequencing, these approaches can be limited in their abilities to distinguish species, due to a lack of differences in ribosomal genes between closely related species and strains. Further, marker gene sequencing cannot be used for downstream analysis such as detecting genetic variation or assembling genomes. Whole metagenome sequencing and marker gene sequencing can therefore function as complementary approaches for interrogating the eukaryotic portion of microbiomes.

## Conclusions

Taken together, the work reported here demonstrates that eukaryotes can be effectively detected from whole metagenome sequencing data with a database of conserved eukaryotic genes. As more metagenomic sequencing data becomes available from host associated and environmental microbiomes, tools like EukDetect will reveal the contributions of microbial eukaryotes to diverse environments.

## Methods

### Identifying universal eukaryotic marker genes in microbial eukaryotes

Microbial eukaryotic genomes were downloaded from NCBI GenBank for all species designated as “Fungi”, “Protists”, “Other”, non-vertebrate metazoans, and non-streptophyte archaeplastids. One genome was downloaded for each species; priority was given to genomes designated as ‘reference genomes’ or ‘representative genomes’. If a species with multiple genomes did not have a designated representative or reference genome, the genome assembly that appeared most contiguous was selected. In addition to the GenBank genomes, 314 genomes and transcriptomes comprised of 282 protists, 30 archaeplastids, and 2 metazoans curated by the EukProt project were downloaded [31].

To identify marker genes in eukaryotic genomes, we ran the Benchmarking Universal Single-Copy Orthologs (BUSCO) version 4 pipeline on all eukaryotic genomes with the Eukaryota OrthoDB version 10 gene models (255 universal eukaryote marker genes) using the augustus optimization parameter (command --long) [30,35]. To ensure that no bacterial sequences were erroneously annotated with this pipeline, we also ran BUSCO with the same parameters on a set of 971 common human gut bacteria from the Culturable Genome Reference [26] and found that this pipeline annotated 41 of the 255 universal marker genes in one or more bacteria. We discarded these predicted markers from each set of BUSCO predicted genes in eukaryotes. While these marker genes comprised ~16% of the total marker genes, removing them from the database reduced the number of simulated reads from the CGR genomes that align to the eukaryotic marker genes from 28,314 to zero. The 214 marker genes never predicted in a bacterial genome are listed in Table S1.

### Constructing the EukDetect database

After running the BUSCO pipeline on each eukaryotic genome and discarding gene models that were annotated in bacterial genomes, we extracted full length nucleotide and protein sequences from each complete BUSCO gene. Protein sequences were used for filtering purposes only. We examined the length distribution of the predicted proteins of each potential marker gene, and genes whose proteins were in the top or bottom 5% of length distributions were discarded. We masked simple repetitive elements with RepeatMasker version open-4.0.7 and discarded genes where 10% or more of the sequence was masked.

To further reduce bacterial contamination, we simulated 2 million sequencing reads from each of 52,515 environmental bacterial metagenome assembled genomes from the GEM catalogue [65] and 4,644 human-associated bacterial metagenome assembled genomes from the UHGG catalogue [66], and aligned these reads to the nucleotide marker gene set. In total, bacterial simulated reads aligned to portions of 8 eukaryotic marker genes. These 8 marker genes were discarded.

In addition to bacterial contamination, eukaryotic genomes can erroneously contain sequences from other eukaryotic genomes [23]. To minimize the impact of this type of contamination, we performed a modified alien index search [67,68] for all marker genes. The protein sequences of all marker genes were aligned to the NCBI nr database using DIAMOND [69]. To calculate the alien index (AI) for a given gene, an in-group lineage is chosen (for example, for the fungus *Saccharomyces cerevisiae*, the phylum Ascomycota may be chosen as the in-group). Then, the best scoring hit for the gene within the group is compared to the best scoring hit outside of the group (for *S. cerevisiae*, to all taxa that are not in the Ascomycota phylum). Any hits to the recipient species itself are skipped. The AI is given by the formula: 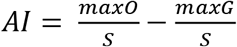 where maxO is the bitscore of the best-scoring hit outside the group lineage, maxG is the bitscore of the best-scoring hit inside the group lineage, and S is the maximum possible bitscore of the gene (i.e., the bitscore of the marker gene aligned to itself). Marker genes where the AI was equal to 1, indicating that there is no in-group hit and the best out-group hit is a perfect match, were discarded.

After filtering, we used CD-HIT version 4.7 to collapse sequences that were greater than 99% identical [70]. For a small number of clusters of genomes, all or most of the annotated BUSCO genes were greater than 99% identical. This arose from errors in taxonomy (in some cases, from fungi with a genome sequence deposited both under the anamorph and teleopmorph name) and from genomes submitted to GenBank with a genus designation only. In these cases, one genome was retained from the collapsed cluster. Some collapsed genes were gene duplicates from the same genome which are designated with the flag “SSCollapse” in the EukDetect database, where “SS” designates “same species”. Genes that were greater than 99% identical between species in the same genus were collapsed, re-annotated as representing NCBI Taxonomy ID associated with the last common ancestor of all species in the collapsed cluster, and annotated with the flag “SGCollapse”. Genes that were collapsed where the species in the collapsed cluster came from multiple NCBI Taxonomy genera were collapsed, annotated as representing the NCBI Taxonomy ID associated with the last common ancestor of all species in the collapsed cluster, and annotated with the flag “MGCollapse”. The EukDetect database is available from Figshare (https://doi.org/10.6084/m9.figshare.12670856.v7). A smaller database of 50 conserved BUSCO genes that could potentially be used for phylogenetics is available from Figshare (https://doi.org/10.6084/m9.figshare.12693164.v2). These 50 marker genes are listed in Table S3.

### The EukDetect pipeline

The EukDetect pipeline uses the Snakemake workflow engine [71]. Sequencing reads are aligned to the EukDetect marker database with Bowtie2 [72]. Reads are filtered for mapping quality greater than 30 and alignments that are less than 80% of the read length of the gene are discarded. Aligned reads are then filtered for sequence complexity with komplexity (https://github.com/eclarke/komplexity) and are discarded below a complexity threshold of 0.5. Duplicate reads are removed with samtools [73]. Reads aligning to marker genes are counted and the percent sequence identity of reads is calculated. The marker genes in the EukDetect database are each linked to NCBI taxonomy IDs. Only taxonomy IDs where more than 4 reads align to 2 or more marker genes are reported in the final results by EukDetect.

Further quality filtering is subsequently performed to reduce false positives arising from various off-target read alignment scenarios (i.e., where sequencing reads originate from one taxon but align to marker genes belonging to another taxon). One cause of off-target alignment is when a marker gene is present in the genome of the off-target species but not in the genome of the source species, due to a gap in the source species’ genome assembly or from the removal of a marker gene by quality control during database construction. A second cause of off-target alignment is when reads originating from one species align spuriously to a marker gene in another species due to sequencing errors or high local similarity between genes.

When more than one species within a genus contains aligned reads passing the read count and marker count filters described above, EukDetect examines the alignments to these species to determine if they could be off-target alignments. First, species are sorted in descending order by maximum number of aligned reads and marker gene bases with at least one aligned read (i.e, marker gene coverage). The species with the highest number of aligned reads and greatest coverage is considered the primary species. In the case of ties, or when the species with the maximum coverage and the maximum number of reads are not the same, all tied species are considered the primary species set. Next, each additional species in the genus is examined to determine whether its aligned sequencing reads could be spurious alignments of reads from any of the species in the primary set. If we do not find evidence that a non-primary species originated from off-target alignments (details below), it is retained and added to the set of primary species.

The first comparison to assess whether a non-primary species could have arisen from off-target alignment is determining whether all of the marker genes with aligned reads in the non-primary species are in the primary species’ marker gene set in the EukDetect database. If not, it is likely that the primary species has those genes but they are spuriously missing from the database due to genome incompleteness or overly conservative filtering in the BUSCO pipeline. In this case, the non-primary species is designated a spurious result and discarded. Next, for shared marker genes, the global percent identity of all aligned reads across the entire gene is determined for both species. If more than half of these shared genes have alignments with an equivalent or greater percent identity in the non-primary compared with the primary species, the non-primary species is considered to be a true positive and added to the set of primary species. Otherwise, it is discarded. In cases where five or fewer genes are shared between the primary and non-primary species, all of the shared genes must have an equivalent or greater percent identity in the non-primary species for it to be retained. This process is performed iteratively across all species within the genus.

Simulations demonstrated that in most cases, this method prevents false positives from single-species simulations and accurately detects mixtures of closely related species (Figure 3C-D). However, in some cases where a species is very close to the detection limit and the other species is present at a high abundance, EukDetect may not detect the low abundance species. EukDetect retains files with alignments for all taxa before any filtering occurs (see Figure S1) for users who wish to examine or differently filter them. We recommend exploring these unfiltered results when a specific low-abundance species is expected in a microbiome or in cases where two very closely related species are thought to be present in a sample at different abundances.

Results are reported both as a count, coverage, and percent sequence identity table for each taxonomy IDs that passes filtering, and as a taxonomy of all reported reads constructed with ete3 [74]. A schematic of the EukDetect pipeline is depicted in Figure S1.

### Simulated reads

Paired-end Illumina reads were simulated from one human-associated fungus (*Candida albicans*) and one soil fungus (*Trichoderma harzianum*), one human-associated protist (*Entameoba dispar*) and one environmental protist (the widely distributed ocean haptophyte *Emiliania huxleyi*), and one human-associated helminth (*Schistosoma mansoni*) and one plant pathogenic nematode (*Globodera rostochiensis*) with InSilicoSeq [75]. Each species was simulated at 19 coverage depths: 0.0001x, 0.001x, 0.01x, 0.02x, 0.03x, 0.04x, 0.05x, 0.1x, 0.25x, 0.5x, 0.75x, 1x, 2x, 3x, 4x, and 5x. Each simulation was repeated 10 times. Simulated reads were processed with the EukDetect pipeline.

### Analysis of public datasets

All sequencing data and associated metadata were taken from public databases and published studies. Sequencing data for determining eukaryotic GI tract distribution was downloaded from the European Nucleotide Archive under accession PRJEB28097 [41,42]. Sequencing data and metadata for the DIABIMMUNE three country cohort was downloaded directly from the DIABIMMUNE project website (https://diabimmune.broadinstitute.org/diabimmune/three-country-cohort) [46]. DNA and RNA sequencing from the IHMP IBDMDB project was downloaded from the NCBI SRA under Bioproject PRJNA398089 [49]. Sequencing data from the *Arabidopsis thaliana* leaf microbiome was downloaded from the European Nucleotide Archive under accession PRJEB31530 [61]. Sequencing data from the Baltic Sea microbiome was downloaded from the NCBI SRA under accessions PRJNA273799 and PRJEB22997 [55,76].

## Declarations

### Ethics approval and consent to participate

Not applicable.

### Consent for publication

Not applicable.

### Availability of data and material

EukDetect is implemented in Python and is available on github (https://github.com/allind/EukDetect). The EukDetect database can be downloaded from Figshare (https://doi.org/10.6084/m9.figshare.12670856.v7).

All datasets analyzed in the current study are publicly available. Sequencing data for determining eukaryotic GI tract distribution was downloaded from the European Nucleotide Archive under accession PRJEB28097 [41,42]. Sequencing data and metadata for the DIABIMMUNE three country cohort was downloaded directly from the DIABIMMUNE project website (https://diabimmune.broadinstitute.org/diabimmune/three-country-cohort) [46]. DNA and RNA sequencing from the IHMP IBDMDB project was downloaded from the NCBI SRA under Bioproject PRJNA398089 [49]. Sequencing data from the *Arabidopsis thaliana* leaf microbiome was downloaded from the European Nucleotide Archive under accession PRJEB31530 [61]. Sequencing data from the Baltic Sea microbiome was downloaded from the NCBI SRA under accessions PRJNA273799 and PRJEB22997 [55,76].

### Authors’ contributions

ALL and KSP designed the study. ALL developed the method and analyzed the data. ALL and KSP interpreted the data and wrote the manuscript.

## Acknowledgements

The authors thank Chunyu Zhao and Jason Shi for valuable contributions to testing.

## Additional information

### Supplementary information

**Additional file 1: Figure S1.** Percentage of simulated bacterial reads from 971 human gut microbiome bacteria that align to 2,449 eukaryotic genomes (see Figure 1A).

**Additional file 2. Figure S2.** (A) Number of aligned reads to a species versus the median coverage of observed marker genes for that species. Median coverage and aligned read counts was calculated for each species within each of the 7 datasets analyzed in this work (see Methods). Red box indicates region depicted in (B).

**Additional file 3. Figure S3**. Schematic of the EukDetect pipeline. (PDF)

**Additional file 4: Figure S4**. Distribution of Blastocystis in the gastrointestinal tract taken from biopsies. Fungi were detected at all sites in the large intestine and terminal ileum, in both lumen and mucosal samples. Slashes indicate no Blastocystis detected in any samples from that site. (PDF)

**Additional file 5: Figure S5.** Distribution of fungal species in the gastrointestinal tract taken from biopsies. Fungi were detected at all sites in the large intestine in both lumen and mucosal samples, and in the lumen of the terminal ileum. One biopsy of gastric antrum mucosa in the stomach contained a Malassezia yeast. Slashes indicate no fungi detected in any samples from that site. (PDF)

**Additional file 6: Figure S6.** (A) Counts of eukaryotic taxa observed in each Baltic Sea sample. (B) Counts of detected chloroplastid subgroups in different environments. (C) Counts of detected stramenopile subgroups in different environments. (D) Counts of detected metazoan subgroups in different environments.

**Additional file 7: Table S1.** OrthoDB v10 Eukaryota marker genes that were not predicted in any of 971 bacterial genomes. (CSV)

**Additional file 8: Table S2.** Genomes with species-level marker genes in the EukDetect database. (CSV)

**Additional file 9: Table S3**. Subset of 50 conserved marker genes found broadly across microbial eukaryotes. (CSV)

**Additional file 10: Table S4.** Eukaryotes in gut biopsy and stool samples. (CSV)

**Additional file 11: Table S5.** Eukaryotes in DIABIMMUNE three-country cohort samples. (CSV)

**Additional file 12: Table S6.** Eukaryotes in IHMP IBD samples. (CSV)

**Additional file 13: Table S7.** Eukaryotes in Baltic Sea samples. (CSV)

**Additional file 14: Table S8.** Eukaryotes from *Arabidopsis thaliana* leaf samples. (CSV)

## References

1. Bik HM, Porazinska DL, Creer S, Caporaso JG, Knight R, Thomas WK. Sequencing our way towards understanding global eukaryotic biodiversity. Trends Ecol Evol. 2012;27:233–43.

2. Rodriguez RJ, Jr JFW, Arnold AE, Redman RS. Fungal endophytes: diversity and functional roles. New Phytol. 2009;182:314–30.

3. Akin DE, Borneman WS. Role of Rumen Fungi in Fiber Degradation. J Dairy Sci. 1990;73:3023–32.

4. Kamoun S, Furzer O, Jones JDG, Judelson HS, Ali GS, Dalio RJD, et al. The Top 10 oomycete pathogens in molecular plant pathology. Mol Plant Pathol. 2015;16:413–34.

5. Haque R. Human Intestinal Parasites. J Health Popul Nutr. 2007;25:387–91.

6. Laforest-Lapointe I, Arrieta M-C. Microbial Eukaryotes: a Missing Link in Gut Microbiome Studies. mSystems. American Society for Microbiology (ASM); 2018;3.

7. Parfrey LW, Walters WA, Lauber CL, Clemente JC, Berg-Lyons D, Teiling C, et al. Communities of microbial eukaryotes in the mammalian gut within the context of environmental eukaryotic diversity. Front Microbiol. 2014;5.

8. Sonnenburg ED, Smits SA, Tikhonov M, Higginbottom SK, Wingreen NS, Sonnenburg JL. Diet-induced extinction in the gut microbiota compounds over generations. Nature. 2016;529:212–5.

9. Caron DA, Alexander H, Allen AE, Archibald JM, Armbrust EV, Bachy C, et al. Probing the evolution, ecology and physiology of marine protists using transcriptomics. Nat Rev Microbiol. Nature Publishing Group; 2017;15:6–20.

10. Brussaard L, de Ruiter PC, Brown GG. Soil biodiversity for agricultural sustainability. Agric Ecosyst Environ. 2007;121:233–44.

11. Hernández-Santos N, Klein BS. Through the Scope Darkly: The Gut Mycobiome Comes into Focus. Cell Host Microbe. Elsevier; 2017;22:728–9.

12. Campo J del, Pons MJ, Herranz M, Wakeman KC, Valle J del, Vermeij MJA, et al. Validation of a universal set of primers to study animal-associated microeukaryotic communities. Environ Microbiol. 2019;21:3855–61.

13. Campo J del, Bass D, Keeling PJ. The eukaryome: Diversity and role of microeukaryotic organisms associated with animal hosts. Funct Ecol. 2020;34:2045–54.

14. Parfrey LW, Walters WA, Knight R. Microbial Eukaryotes in the Human Microbiome: Ecology, Evolution, and Future Directions. Front Microbiol. 2011;2:153.

15. Andersen LO, Vedel Nielsen H, Stensvold CR. Waiting for the human intestinal Eukaryotome. ISME J. Nature Publishing Group; 2013;7:1253–5.

16. Lücking R, Aime MC, Robbertse B, Miller AN, Ariyawansa HA, Aoki T, et al. Unambiguous identification of fungi: where do we stand and how accurate and precise is fungal DNA barcoding? IMA Fungus. 2020;11:14.

17. Schoch CL, Seifert KA, Huhndorf S, Robert V, Spouge JL, Levesque CA, et al. Nuclear ribosomal internal transcribed spacer (ITS) region as a universal DNA barcode marker for Fungi. Proc Natl Acad Sci. National Academy of Sciences; 2012;109:6241–6.

18. Knight R, Vrbanac A, Taylor BC, Aksenov A, Callewaert C, Debelius J, et al. Best practices for analysing microbiomes. Nat Rev Microbiol. Nature Publishing Group; 2018;16:410–22.

19. Qin J, Li R, Raes J, Arumugam M, Burgdorf KS, Manichanh C, et al. A human gut microbial gene catalogue established by metagenomic sequencing. Nature. 2010;464:59–65.

20. Nash AK, Auchtung TA, Wong MC, Smith DP, Gesell JR, Ross MC, et al. The gut mycobiome of the Human Microbiome Project healthy cohort. Microbiome. BioMed Central; 2017;5:153.

21. Chehoud C, Albenberg LG, Judge C, Hoffmann C, Grunberg S, Bittinger K, et al. A Fungal Signature in the Gut Microbiota of Pediatric Patients with Inflammatory Bowel Disease. Inflamm Bowel Dis. 2015;21:1948–56.

22. Lu J, Salzberg SL. Removing contaminants from databases of draft genomes. PLOS Comput Biol. 2018;14:e1006277.

23. Steinegger M, Salzberg SL. Terminating contamination: large-scale search identifies more than 2,000,000 contaminated entries in GenBank. Genome Biol. 2020;21:115.

24. Truong DT, Franzosa EA, Tickle TL, Scholz M, Weingart G, Pasolli E, et al. MetaPhlAn2 for enhanced metagenomic taxonomic profiling. Nat Methods. Nature Publishing Group; 2015;12:902–3.

25. Beghini F, McIver LJ, Blanco-Míguez A, Dubois L, Asnicar F, Maharjan S, et al. Integrating taxonomic, functional, and strain-level profiling of diverse microbial communities with bioBakery 3. bioRxiv. Cold Spring Harbor Laboratory; 2020;2020.11.19.388223.

26. Zou Y, Xue W, Luo G, Deng Z, Qin P, Guo R, et al. 1,520 reference genomes from cultivated human gut bacteria enable functional microbiome analyses. Nat Biotechnol. 2019;37:179–85.

27. Marcelino VR, Clausen PTLC, Buchmann JP, Wille M, Iredell JR, Meyer W, et al. CCMetagen: comprehensive and accurate identification of eukaryotes and prokaryotes in metagenomic data. Genome Biol. 2020;21:103.

28. Gentekaki E, Curtis BA, Stairs CW, Klimeš V, Eliáš M, Salas-Leiva DE, et al. Extreme genome diversity in the hyper-prevalent parasitic eukaryote Blastocystis. PLOS Biol. 2017;15:e2003769.

29. McCarthy CGP, Fitzpatrick DA. Pan-genome analyses of model fungal species. Microb Genomics [Internet]. 2019 [cited 2020 Jul 6];5. Available from: https://www.ncbi.nlm.nih.gov/pmc/articles/PMC6421352/

30. Simão FA, Waterhouse RM, Ioannidis P, Kriventseva EV, Zdobnov EM. BUSCO: Assessing genome assembly and annotation completeness with single-copy orthologs. Bioinformatics. 2015;31:3210–2.

31. Richter DJ, Berney C, Strassert JFH, Burki F, Vargas C de. EukProt: a database of genome-scale predicted proteins across the diversity of eukaryotic life. bioRxiv. Cold Spring Harbor Laboratory; 2020;2020.06.30.180687.

32. Saary P, Mitchell AL, Finn RD. Estimating the quality of eukaryotic genomes recovered from metagenomic analysis with EukCC. Genome Biol. 2020;21:244.

33. Wu D, Jospin G, Eisen JA. Systematic Identification of Gene Families for Use as “Markers” for Phylogenetic and Phylogeny-Driven Ecological Studies of Bacteria and Archaea and Their Major Subgroups. PLOS ONE. Public Library of Science; 2013;8:e77033.

34. Waterhouse RM, Seppey M, Simão FA, Manni M, Ioannidis P, Klioutchnikov G, et al. BUSCO Applications from Quality Assessments to Gene Prediction and Phylogenomics. Mol Biol Evol. 2018;35:543–8.

35. Kriventseva EV, Kuznetsov D, Tegenfeldt F, Manni M, Dias R, Simão FA, et al. OrthoDB v10: sampling the diversity of animal, plant, fungal, protist, bacterial and viral genomes for evolutionary and functional annotations of orthologs. Nucleic Acids Res. 2019;47:D807–11.

36. McCutcheon JP, Moran NA. Extreme genome reduction in symbiotic bacteria. Nat Rev Microbiol. Nature Publishing Group; 2012;10:13–26.

37. Stukenbrock EH. The Role of Hybridization in the Evolution and Emergence of New Fungal Plant Pathogens. Phytopathology. 2016;106:104–12.

38. Marcet-Houben M, Gabaldón T. Beyond the Whole-Genome Duplication: Phylogenetic Evidence for an Ancient Interspecies Hybridization in the Baker’s Yeast Lineage. PLOS Biol. Public Library of Science; 2015;13:e1002220.

39. Jain C, Rodriguez-R LM, Phillippy AM, Konstantinidis KT, Aluru S. High throughput ANI analysis of 90K prokaryotic genomes reveals clear species boundaries. Nat Commun. Nature Publishing Group; 2018;9:5114.

40. Tropini C, Earle KA, Huang KC, Sonnenburg JL. The gut microbiome: Connecting spatial organization to function. Cell Host Microbe. 2017;21:433–42.

41. Suez J, Zmora N, Zilberman-Schapira G, Mor U, Dori-Bachash M, Bashiardes S, et al. Post-Antibiotic Gut Mucosal Microbiome Reconstitution Is Impaired by Probiotics and Improved by Autologous FMT. Cell. 2018;174:1406–1423.e16.

42. Zmora N, Zilberman-Schapira G, Suez J, Mor U, Dori-Bachash M, Bashiardes S, et al. Personalized Gut Mucosal Colonization Resistance to Empiric Probiotics Is Associated with Unique Host and Microbiome Features. Cell. 2018;174:1388–1405.e21.

43. Suhr MJ, Hallen-Adams HE. The human gut mycobiome: pitfalls and potentials--a mycologists perspective. Mycologia. 2015;107:1057–73.

44. Moe KT, Singh M, Howe J, Ho LC, Tan SW, Chen XQ, et al. Experimental Blastocystis hominis infection in laboratory mice. Parasitol Res. 1997;83:319–25.

45. Bäckhed F, Roswall J, Peng Y, Feng Q, Jia H, Kovatcheva-Datchary P, et al. Dynamics and Stabilization of the Human Gut Microbiome during the First Year of Life. Cell Host Microbe. 2015;17:690–703.

46. Vatanen T, Kostic AD, d’Hennezel E, Siljander H, Franzosa EA, Yassour M, et al. Variation in Microbiome LPS Immunogenicity Contributes to Autoimmunity in Humans. Cell. Elsevier; 2016;165:842–53.

47. Olm MR, West PT, Brooks B, Firek BA, Baker R, Morowitz MJ, et al. Genome-resolved metagenomics of eukaryotic populations during early colonization of premature infants and in hospital rooms. Microbiome [Internet]. 2019 [cited 2020 Jul 6];7. Available from: https://www.ncbi.nlm.nih.gov/pmc/articles/PMC6377789/

48. Fujimura KE, Sitarik AR, Havstad S, Lin DL, Levan S, Fadrosh D, et al. Neonatal gut microbiota associates with childhood multisensitized atopy and T cell differentiation. Nat Med. 2016;22:1187–91.

49. Lloyd-Price J, Arze C, Ananthakrishnan AN, Schirmer M, Avila-Pacheco J, Poon TW, et al. Multi-omics of the gut microbial ecosystem in inflammatory bowel diseases. Nature. Nature Publishing Group; 2019;569:655–62.

50. Scanlan PD, Stensvold CR, Rajilić-Stojanović M, Heilig HGHJ, De Vos WM, O’Toole PW, et al. The microbial eukaryote Blastocystis is a prevalent and diverse member of the healthy human gut microbiota. FEMS Microbiol Ecol. 2014;90:326–30.

51. Limon JJ, Tang J, Li D, Wolf AJ, Michelsen KS, Funari V, et al. Malassezia Is Associated with Crohn’s Disease and Exacerbates Colitis in Mouse Models. Cell Host Microbe. 2019;25:377–388.e6.

52. Sokol H, Leducq V, Aschard H, Pham H-P, Jegou S, Landman C, et al. Fungal microbiota dysbiosis in IBD. Gut. 2017;66:1039–48.

53. Aykut B, Pushalkar S, Chen R, Li Q, Abengozar R, Kim JI, et al. The fungal mycobiome promotes pancreatic oncogenesis via activation of MBL. Nature. 2019;574:264–7.

54. Leppäranta M, Myrberg K. Physical Oceanography of the Baltic Sea. Springer Science & Business Media; 2009.

55. Hugerth LW, Larsson J, Alneberg J, Lindh MV, Legrand C, Pinhassi J, et al. Metagenome-assembled genomes uncover a global brackish microbiome. Genome Biol. 2015;16:279.

56. Wylezich C, Karpov SA, Mylnikov AP, Anderson R, Jürgens K. Ecologically relevant choanoflagellates collected from hypoxic water masses of the Baltic Sea have untypical mitochondrial cristae. BMC Microbiol. 2012;12:271.

57. Radwan O, Gunasekera TS, Ruiz ON. Draft Genome Sequence of Lecanicillium sp. Isolate LEC01, a Fungus Capable of Hydrocarbon Degradation. Microbiol Resour Announc [Internet]. American Society for Microbiology; 2019 [cited 2020 Dec 14];8. Available from: https://mra.asm.org/content/8/15/e01744-18

58. Thureborn P, Lundin D, Plathan J, Poole AM, Sjöberg B-M, Sjöling S. A metagenomics transect into the deepest point of the Baltic Sea reveals clear stratification of microbial functional capacities. PloS One. 2013;8:e74983.

59. Sjöqvist C, Godhe A, Jonsson PR, Sundqvist L, Kremp A. Local adaptation and oceanographic connectivity patterns explain genetic differentiation of a marine diatom across the North Sea-Baltic Sea salinity gradient. Mol Ecol. 2015;24:2871–85.

60. Jacob S, Legrand D, Chaine AS, Bonte D, Schtickzelle N, Huet M, et al. Gene flow favours local adaptation under habitat choice in ciliate microcosms. Nat Ecol Evol. Nature Publishing Group; 2017;1:1407–10.

61. Regalado J, Lundberg DS, Deusch O, Kersten S, Karasov T, Poersch K, et al. Combining whole-genome shotgun sequencing and rRNA gene amplicon analyses to improve detection of microbe-microbe interaction networks in plant leaves. ISME J. Nature Publishing Group; 2020;1–15.

62. Osman M, El Safadi D, Cian A, Benamrouz S, Nourrisson C, Poirier P, et al. Prevalence and Risk Factors for Intestinal Protozoan Infections with Cryptosporidium, Giardia, Blastocystis and Dientamoeba among Schoolchildren in Tripoli, Lebanon. PLoS Negl Trop Dis. 2016;10:e0004496.

63. Ahrendt SR, Quandt CA, Ciobanu D, Clum A, Salamov A, Andreopoulos B, et al. Leveraging single-cell genomics to expand the fungal tree of life. Nat Microbiol. Nature Publishing Group; 2018;3:1417–28.

64. West PT, Probst AJ, Grigoriev IV, Thomas BC, Banfield JF. Genome-reconstruction for eukaryotes from complex natural microbial communities. Genome Res. Cold Spring Harbor Laboratory Press; 2018;28:569–80.

65. Nayfach S, Roux S, Seshadri R, Udwary D, Varghese N, Schulz F, et al. A genomic catalog of Earth’s microbiomes. Nat Biotechnol. Nature Publishing Group; 2020;1–11.

66. Almeida A, Nayfach S, Boland M, Strozzi F, Beracochea M, Shi ZJ, et al. A unified catalog of 204,938 reference genomes from the human gut microbiome. Nat Biotechnol. Nature Publishing Group; 2021;39:105–14.

67. Wisecaver JH, Alexander WG, King SB, Todd Hittinger C, Rokas A. Dynamic Evolution of Nitric Oxide Detoxifying Flavohemoglobins, a Family of Single-Protein Metabolic Modules in Bacteria and Eukaryotes. Mol Biol Evol. Oxford University Press; 2016;33:1979–87.

68. Gladyshev EA, Meselson M, Arkhipova IR. Massive horizontal gene transfer in bdelloid rotifers. Science. 2008;320:1210–3.

69. Buchfink B, Xie C, Huson DH. Fast and sensitive protein alignment using DIAMOND. Nat Methods. Nature Publishing Group; 2015;12:59–60.

70. Fu L, Niu B, Zhu Z, Wu S, Li W. CD-HIT: accelerated for clustering the next-generation sequencing data. Bioinforma Oxf Engl. 2012;28:3150–2.

71. Köster J, Rahmann S. Snakemake—a scalable bioinformatics workflow engine. Bioinformatics. Oxford Academic; 2012;28:2520–2.

72. Langmead B, Salzberg SL. Fast gapped-read alignment with Bowtie 2. Nat Methods. 2012;9:357–9.

73. Li H, Handsaker B, Wysoker A, Fennell T, Ruan J, Homer N, et al. The Sequence Alignment/Map format and SAMtools. Bioinforma Oxf Engl. 2009;25:2078–9.

74. Huerta-Cepas J, Serra F, Bork P. ETE 3: Reconstruction, Analysis, and Visualization of Phylogenomic Data. Mol Biol Evol. Oxford Academic; 2016;33:1635–8.

75. Gourlé H, Karlsson-Lindsjö O, Hayer J, Bongcam-Rudloff E. Simulating Illumina metagenomic data with InSilicoSeq. Bioinforma Oxf Engl. 2019;35:521–2.

76. Alneberg J, Sundh J, Bennke C, Beier S, Lundin D, Hugerth LW, et al. BARM and BalticMicrobeDB, a reference metagenome and interface to meta-omic data for the Baltic Sea. Sci Data. Nature Publishing Group; 2018;5:180146.

