## Supplementary figures and images for "Accurate and sensitive detection of microbial eukaryotes from whole metagenome shotgun sequencing"

### Figure S1

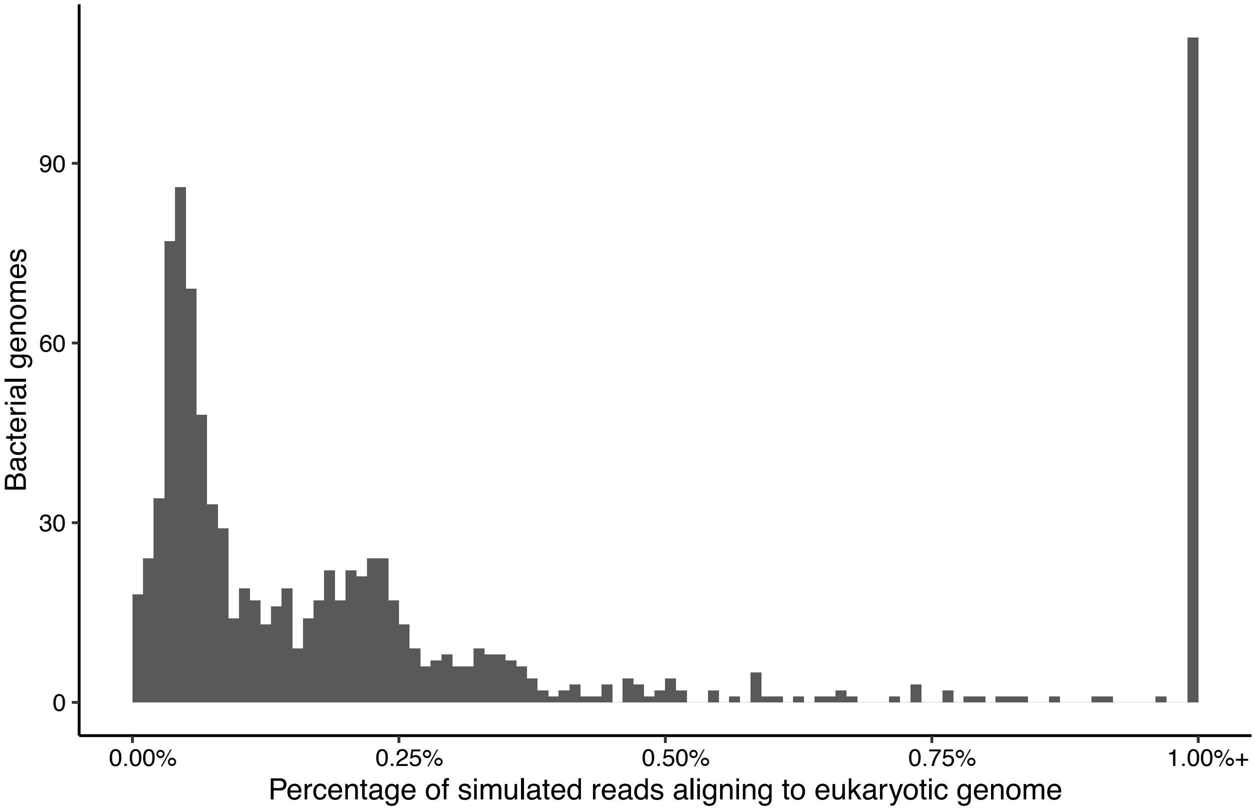

### Figure S2

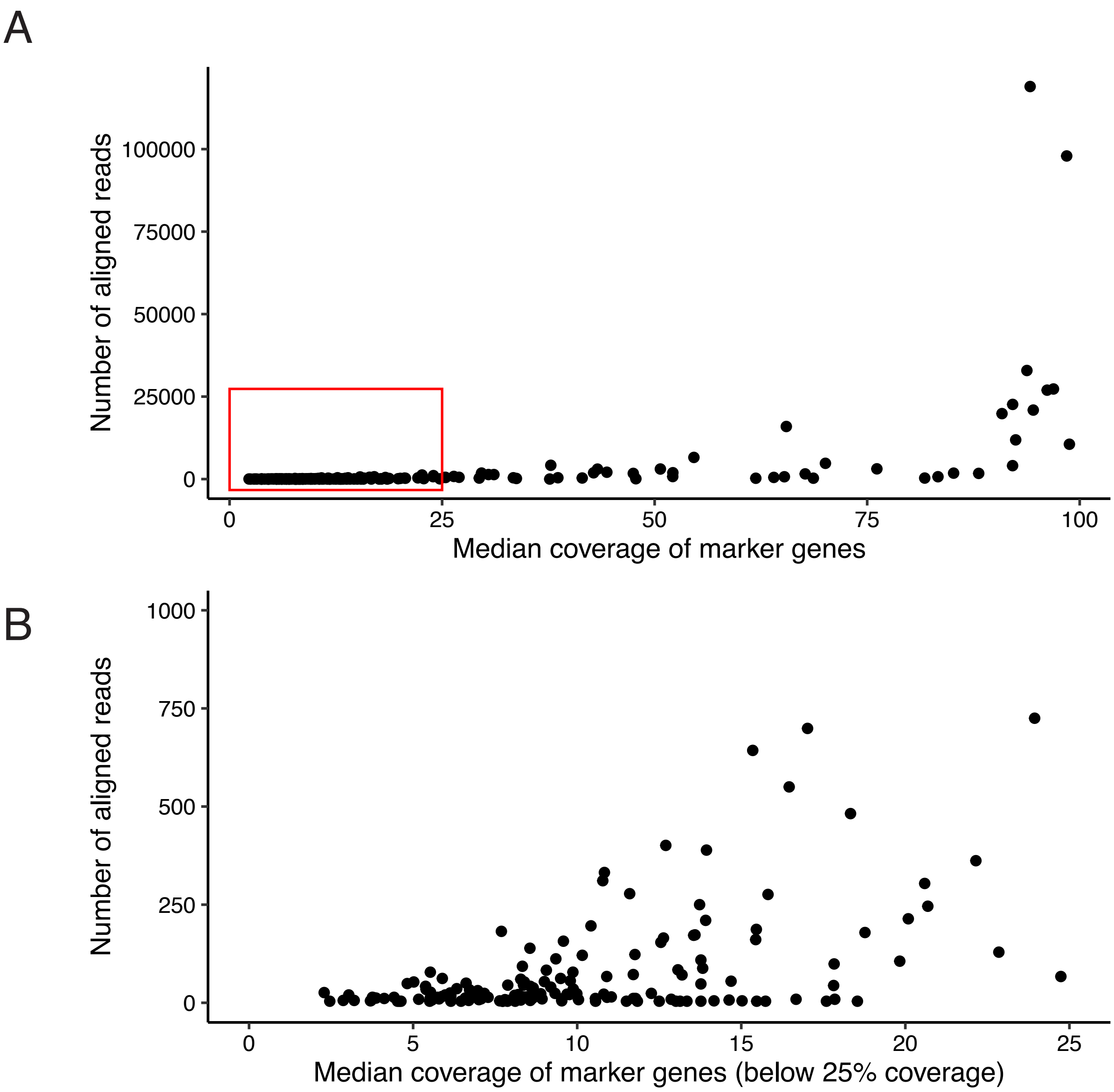

### Figure S3

Figure S1. Schematic of the EukDetect pipeline.

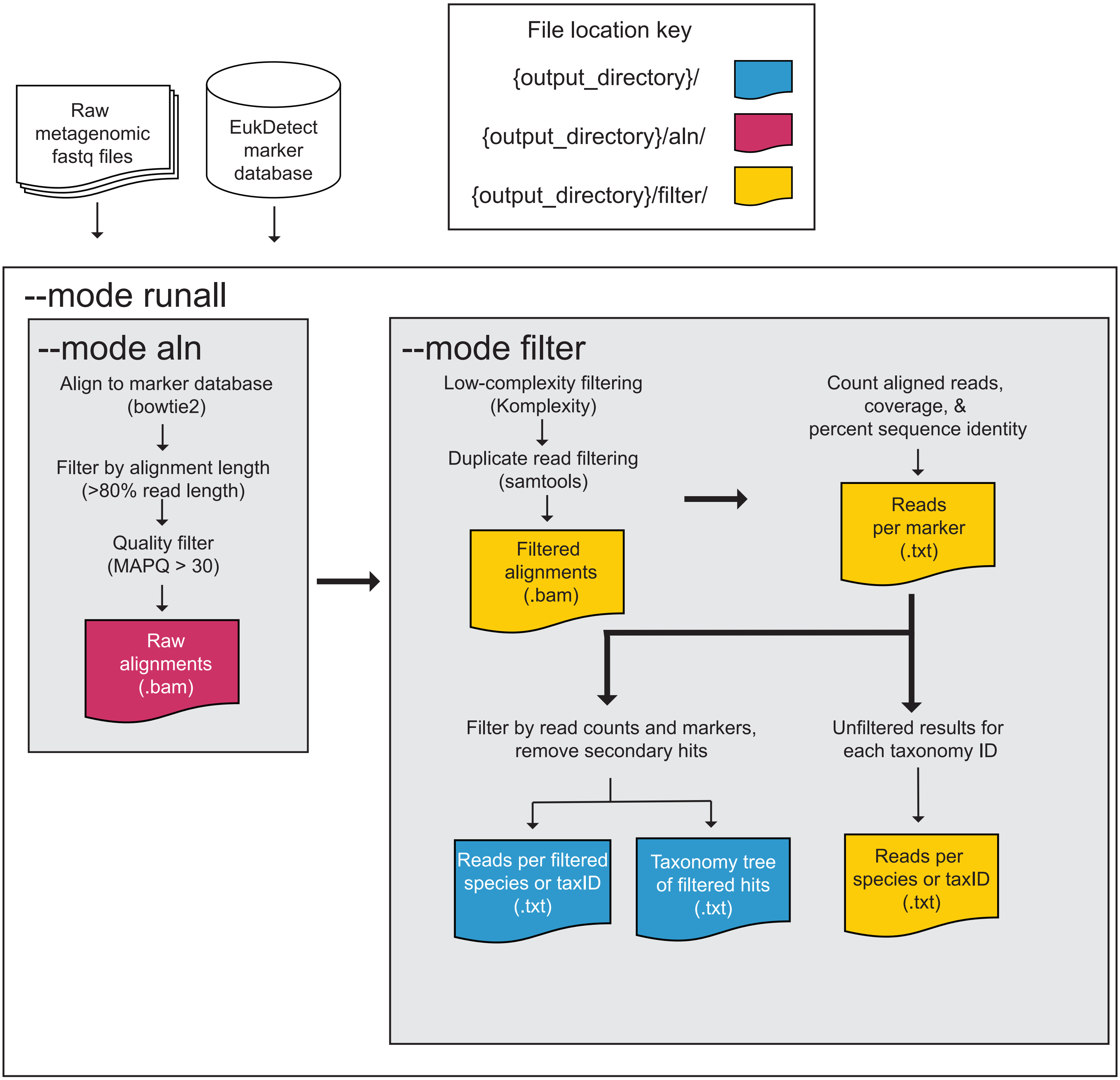

### Figure S6

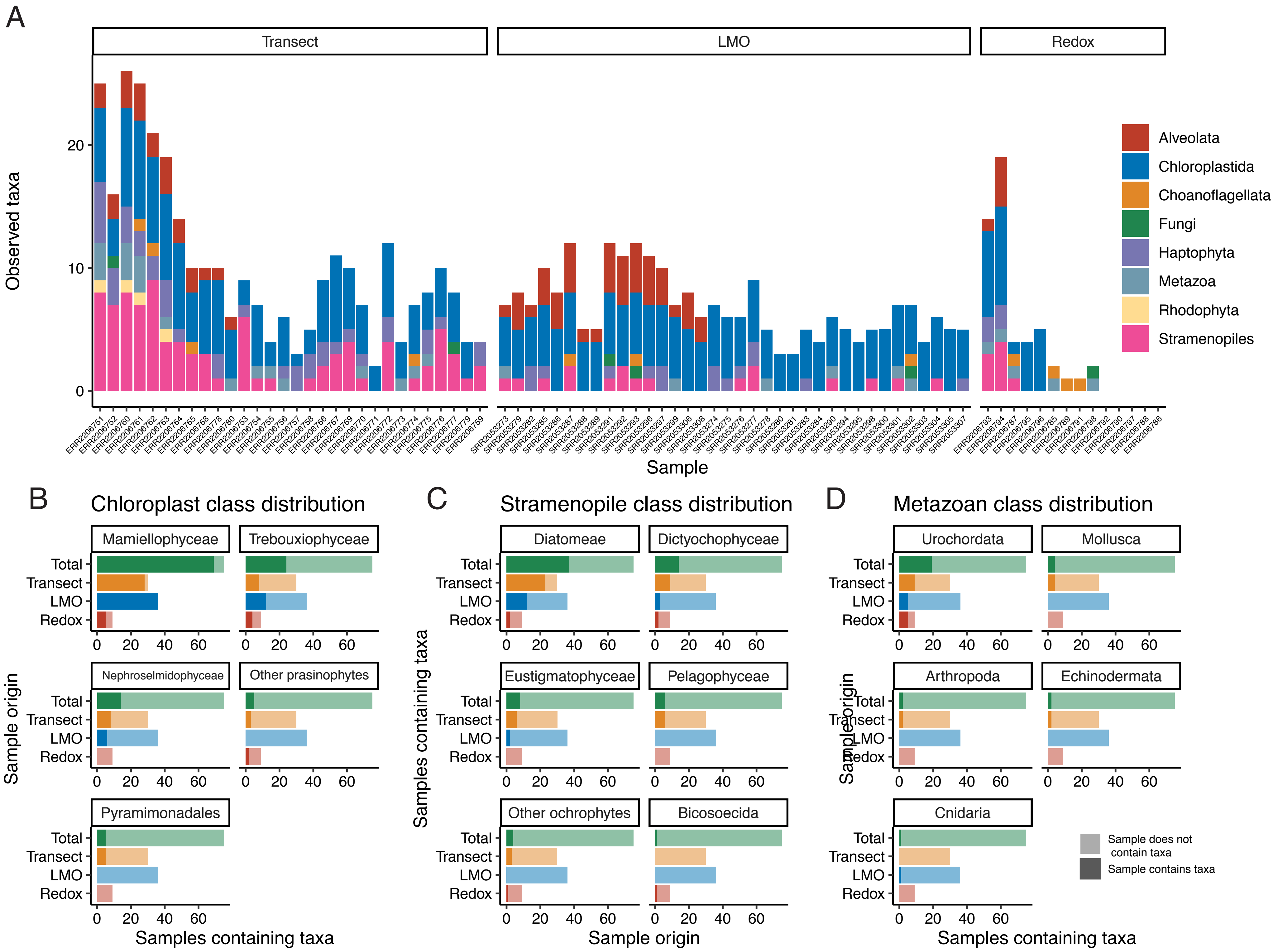
