## Supplementary material for "Accurate and sensitive detection of microbial eukaryotes from whole metagenome shotgun sequencing": Figure S4

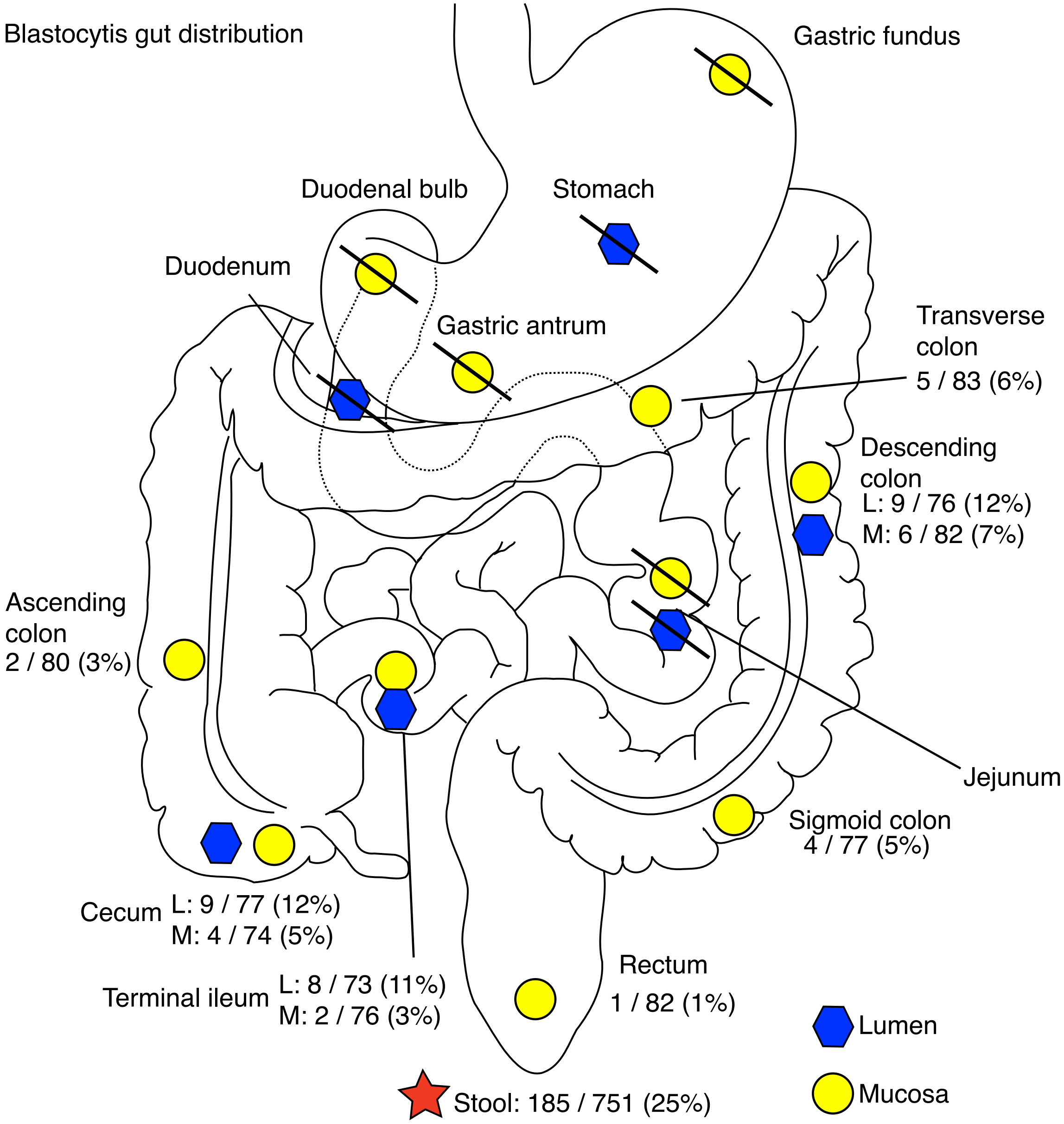


Figure S2. Distribution of *Blastocystis* in the gastrointestinal tract taken from biopsies. Fungi were detected at all sites in the large intestine and terminal ileum, in both lumen and mucosal samples. Slashes indicate no *Blastocystis* detected in any samples from that site.
