## Supplementary material for "Accurate and sensitive detection of microbial eukaryotes from whole metagenome shotgun sequencing": Figure S5

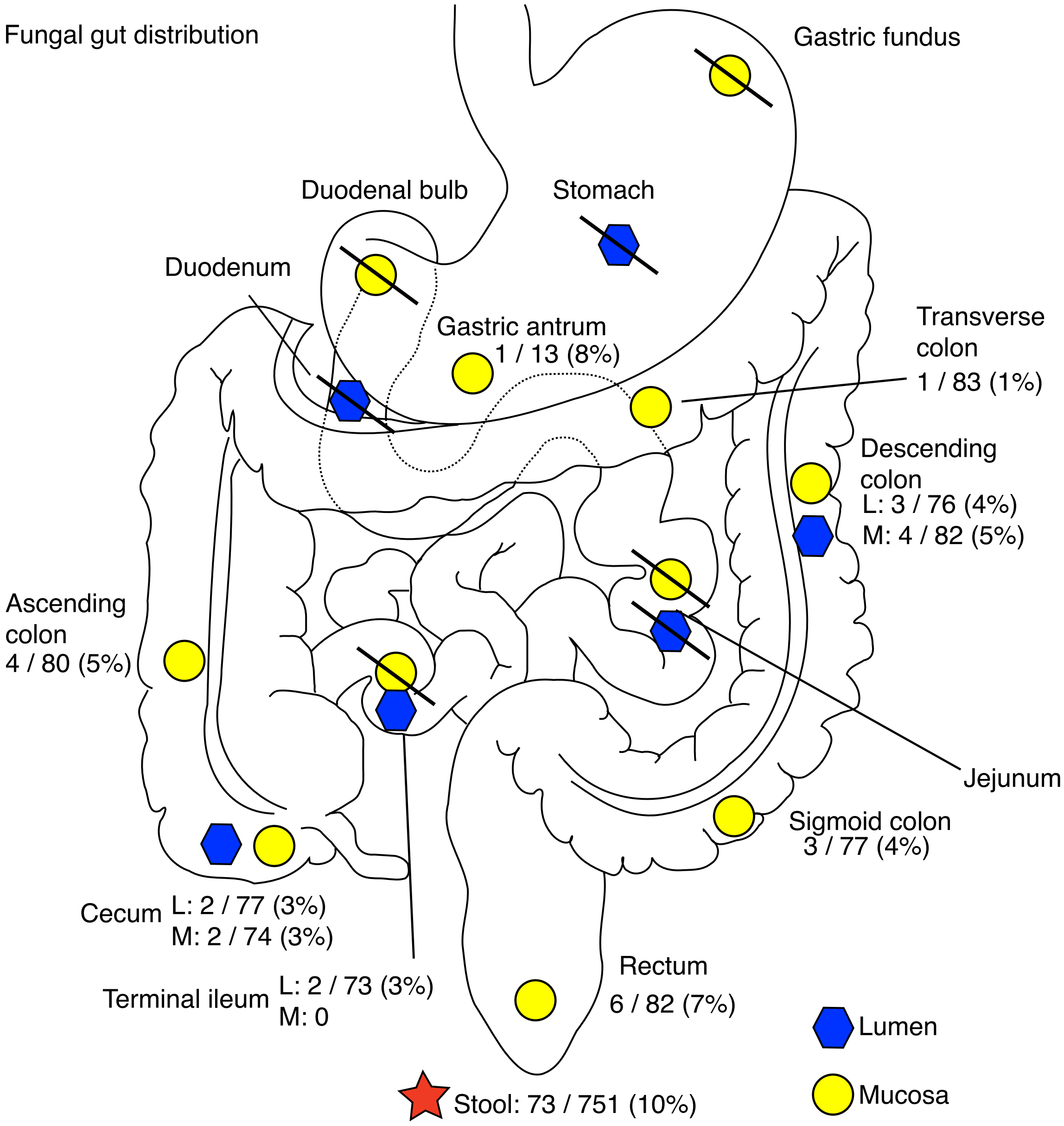


Figure S3. Distribution of fungal species in the gastrointestinal tract taken from biopsies. Fungi were detected at all sites in the large intestine in both lumen and mucosal samples, and in the lumen of the terminal ileum. One biopsy of gastric antrum mucosa in the stomach contained a *Malassezia* yeast. Slashes indicate no fungi detected in any samples from that site.
